# WNT11 is a novel ligand for ROR2 in human breast cancer

**DOI:** 10.1101/2020.12.18.423402

**Authors:** Kerstin Menck, Saskia Heinrichs, Darius Wlochowitz, Maren Sitte, Helen Noeding, Andreas Janshoff, Hannes Treiber, Torben Ruhwedel, Bawarjan Schatlo, Christian von der Brelie, Stefan Wiemann, Tobias Pukrop, Tim Beißbarth, Claudia Binder, Annalen Bleckmann

**Affiliations:** Dept. of Medicine A, Hematology, Oncology, and Pneumology, University Hospital Münster, Münster, Germany; West German Cancer Center, University Hospital Münster, Münster, Germany; Dept. of Hematology/Medical Oncology, University Medical Center Göttingen, Göttingen, Germany; Dept. of Medical Bioinformatics, University Medical Center Göttingen, Göttingen, Germany; Institute for Physical Chemistry, Georg August University Göttingen, Göttingen, Germany; Dept. of Neurogenetics, Max Planck Institute of Experimental Medicine, Göttingen, Germany; Dept. of Neurosurgery, University Medical Center Göttingen, Göttingen, Germany; Division of Molecular Genome Analysis, German Cancer Research Center, Heidelberg, Germany; Dept. of Internal Medicine III, Hematology and Medical Oncology, University Hospital Regensburg, Regensburg, Germany

**Keywords:** Breast cancer metastasis, ROR2, WNT11, BRCAness, network analysis

## Abstract

Breast cancer has been associated with activation of the WNT signaling pathway, although the underlying molecular mechanisms are still unclear. Here, we found the WNT receptor ROR2 to be highly expressed in aggressive breast tumors and associated with worse metastasis-free survival. In order to understand the molecular basis of these observations, we overexpressed ROR2 in human breast cancer cell lines, inducing a BRCAness-like phenotype and rendering them resistant to PARP inhibition. High levels of ROR2 were associated with defects in cell morphology and cell-cell-contacts leading to increased tumor invasiveness. Using gene expression analysis we demonstrated an upregulation of several non-canonical WNT ligands in ROR2-overexpressing breast cancer cells, in particular WNT11. Co-immunoprecipitation confirmed that WNT11 is indeed a novel ligand for ROR2 that interacts with its cysteine-rich domain and triggers the invasion-promoting signaling via RHO/ROCK. Knockdown of WNT11 reversed the pro-invasive phenotype and the cellular changes in ROR2-overexpressing cells. Taken together, our studies revealed a novel auto-stimulatory loop in which ROR2 triggers the expression of its own ligand, WNT11, resulting in enhanced tumor invasion associated with breast cancer metastasis.

## INTRODUCTION

Breast cancer is the most common cancer in women with more than two million new cases diagnosed in 2018^1^. Patients frequently develop metastases in the course of their disease which limit survival due to the lack of a curative treatment. One signaling pathway that is frequently involved in cancer initiation and progression is the WNT pathway. In mammals it comprises nineteen secreted WNT ligands that can interact with ten different Frizzled (FZD) receptors and various co-receptors^2^. WNT ligands activate different intracellular signaling cascades depending on the specific combination of locally available ligands, receptors and co-receptors^3,4^.

Binding of a canonical WNT ligand (e.g. WNT3a) to a FZD receptor and LRP5/6 co-receptor activates β-catenin-dependent, canonical signaling that results in the expression of WNT-responsive target genes^2^. Other WNT ligands such as WNT5A/B, or WNT11 can bind FZDs and alternative co-receptors (e.g. ROR1/2, RYK, PTK7) and trigger a multitude of β-catenin-independent, non-canonical WNT signaling cascades, the best-studied ones being WNT/Calcium (Ca^2+^) and WNT/planar cell polarity (PCP) signaling. Activation of the WNT/Ca^2+^ pathway is characterized by an increase in intracellular calcium levels that triggers the activation of PKC, NFκB and CREB^5^. In contrast, in WNT/PCP signaling binding of a WNT ligand induces the recruitment of a DVL/RHO/DAAM1 complex that is required for subsequent activation of the RHO/ROCK pathway^6^. In parallel, DVL can activate the small GTPase RAC and downstream JNK signaling (JNK)^7^. Non-canonical WNT signal transduction mostly results in changes in the cytoskeleton, cell motility and morphology^2^.

While aberrant WNT signaling is a hallmark of colorectal cancer, its role in breast cancer is less clear. No driver mutations in typical WNT genes have been detected so far. Nonetheless, several studies point to hyperactive and dysbalanced WNT signaling^8,9^. Especially in basal-like breast cancer, the most unfavorable clinical subtype with early metastasis formation, active non-canonical WNT signaling has been identified and linked to the aggressive behaviour of these breast cancer cells. The highly motile and invasive phenotype of the cancer cells has been mainly attributed to the high expression of WNT5A/B and receptor tyrosine kinase-like orphan receptors 1 and 2 (ROR1/2)^10–13^. In contrast to ROR1, ROR2 was not only found to be highly expressed in basal-like but in 87% of all breast cancers, and high levels were associated with shorter overall survival^14,15^. A significant role of ROR2 in tumor development has already been confirmed *in vivo* in a basal-like TP53-null mouse model of breast cancer where knockdown of ROR2 significantly impaired tumor growth^16^. Apart from the primary tumor tissue, high levels of ROR2 were detectable in lymph node and brain metastases^10,15^, thus suggesting its involvement in tumor progression and metastasis.

ROR2 has been shown to interact with other WNT signaling proteins such as VANGL2^17^ or the WNT co-receptor PTK7^18^. The former is a core component of WNT/PCP signaling which has recently been established as an important factor determining tumor growth and survival in basal-like breast cancer patients^19^. ROR2 harbors a cysteine-rich domain (CRD) in its extracellular part that resembles the WNT protein binding domain of FZD receptors. Indeed, ROR2 has been shown to interact with WNT5A and WNT3A^4,20^, however, the latter failed to activate ROR2-induced signaling^21^. Considering the ambiguous role of WNT5A which has been described to act both as an oncogene as well as a tumor suppressor in breast cancer^22^, it is still unclear whether other WNT ligands exist that can activate ROR2 signaling.

In this study we addressed this question and demonstrated for the first time that WNT11 is a novel ligand for ROR2 in humans. WNT11 binds to the CRD of ROR2 and mediates WNT/PCP signaling via the RHO/ROCK pathway that confers an aggressive phenotype to breast cancer cells. ROR2 and WNT11 are both highly expressed in human brain metastases and linked with short patient survival.

## METHODS

### Cell lines, transfections and viability assays

MCF-7, T-47D, MDA-MB-231 and SK-BR-3 cells (DSMZ, ATCC) were cultured in RPMI-1640, BT-474 cells in DMEM/F12, all supplemented with 10% heat-inactivated (56°C, 30 min) fetal calf serum (FCS). For siRNA-mediated gene knockdown, cells were transfected with 10 nM siRNA (santa cruz) using the RNAimax transfection reagent (Invitrogen). Cells were used 24 h post transfection for functional studies and 72h post transfection for expression analysis. To achieve gene overexpression, cells were transfected with Fugene HD transfection reagent (Promega) and stable clones generated by geneticin or zeocin selection (750 µg/ml or 100 µg/ml, respectively). Cell viability upon treatment with olaparib (Selleck chemicals) for 96 h was measured by MTT assay using standard protocols.

### Vectors

The vector for V5-tagged active WNT11 was a gift from Xi He (Addgene plasmid #43824; http://n2t.net/addgene:43824; RRID:Addgene_43824)^23^. The pcDNA3.1/Zeo(+) empty vector was obtained from Invitrogen. The pROR2 vector was kindly provided by Alexandra Schambony. N-terminal ROR2 deletion constructs were generated by PCR-based cloning. They consisted of amino acids 146-943 for ROR2-Δ, 304-943 for ROR2-ΔΔ and 395-943 for ROR2-ΔΔΔ, and were inserted into the empty vector by NheI and XhoI restriction. C-terminal truncation was achieved by creating premature stop codons at amino acid position 467 (ΔPRD) and 783 (ΔPRDΔTKD) using site-directed mutagenesis. Successful cloning was confirmed by sequencing.

### Patient samples and pathway enrichment analysis

Samples of brain metastases were collected from patients previously diagnosed with primary breast cancer during neurosurgical removal. All samples were obtained with informed consent as approved by the local ethics committee (24/10/05). Expression of *ROR1, ROR2, PTK7* and *RYK* in normal and cancerous breast was analysed on microarray data with the IST Online™ database (http://www.medisapiens.com/)^24^ as well as on RNA-Seq data from matched samples of normal and invasive breast carcinoma tissue using the TNMplot database (https://www.tnmplot.com/)^25^. Gene expression data of primary breast cancer patients were compiled from ten public microarray datasets measured on Affymetrix Human Genome HG-U133 Plus 2.0 and HG-U133A arrays as previously described^12^. For RNA-Seq analysis of breast cancer brain metastases (available as GSE161865 on the GEO repository) total RNA was isolated with the Trizol reagent from fresh frozen tissue and RNA integrity was checked with the Bioanalyzer 2100 (Agilent Technologies). For cDNA library preparation the TruSeq Stranded Total RNA sample preparation kit (Illumina) was used, and accurate quantification was performed with the QuantiFluor dsDNA system (Promega). The size range of the generated cDNA libraries was measured with the DNA 1000 chip on the Bioanalyzer 2100 (280 bp). Amplification and sequencing of cDNA libraries were performed using the cBot and HiSeq 2000 (Illumina, SR, 1×51 bp, 8-9 gigabases, >40 mio reads per sample). Sequence images were transformed with Illumina software BaseCaller to bcl-files, which were demultiplexed to fastq-files with CASAVA (v1.8.2). Quality check was done via FastQC (v0.10.1, Babraham Bioinformatics).

Sequence reads were aligned with the reference genome GRCh37 using the STAR RNA-Seq alignment tool^26^, while incorporating database information from Ensembl (v37.73) during the reference indexing step. Gene-level abundances were estimated using the RNA-seq by expectation-maximization (RSEM) algorithm^27^. The relation between gene expression and patient prognosis based on overall survival (OS) was studied among a patient cohort of 32 patients who developed breast cancer brain metastases. One patient sample was discarded from the study after quality control due to small initial RNA-Seq library size, leaving 31 patient samples for further processing and analysis. RSEM estimated counts of patient samples were converted to log2-transformed normalized counts as transcripts per million (TPM). Rank-based enrichment testing was performed for three different WNT pathway signatures using the Wilcoxon rank-sum test, thereby obtaining *P-*values across patient samples per pathway^12^. Transformed −log10(*P-*values) were then subjected to complete-linkage hierarchical clustering based on Pearson correlation as the distance measure. Patient groups in terms of high and low pathway enrichment were obtained using the *cutree* function from the *dendextend* R package^28^.

To conduct Kaplan–Meier survival analysis of gene expression levels of the two-gene signature (ROR2 and WNT11), high/low groups were defined based on the averaged gene expression levels of the gene signature by applying an optimal cutoff value that was computed using the *surv_cutpoint* function from the *survminer* R package (v0.4.8, https://CRAN.R-project.org/package=survminer). Kaplan–Meier survival analysis were performed to assess different survival rates between the groups using the Log-Rank test. Significance p values and hazard ratios (HR) were obtained using the *survival* R package.

### PAM50 molecular subtype classification and BRCAness-like gene signature scoring

The RNA-seq raw data of MCF-7 cells overexpressing ROR2 and of MCF-7 pcDNA/pROR2 cells with control and WNT11 siRNA are available as GSE74383 and GSE161864 on the GEO repository and were pre-processed as described in^12^. RSEM estimated counts of samples were filtered and genes with count per million (CPM)>1 in more than three of the samples were kept using the *edgeR* R package^29^. The gene-level count matrix was then normalized using the “Trimmed Mean of M-values” (TMM) method from the edgeR R package and log2-transformed normalized pseudo counts were obtained for PAM50 subtype classification. Count values were first averaged over triplicates per condition for each gene in the matrix and probability percentage values were estimated for each condition to belong to one of the five molecular subtypes using the *molecular*.*subtyping* function from the *genefu* R Package. To quantify the relevance of a BRCAness-like phenotype, the unsupervised gene set enrichment method called “Gene Set Variation Analysis” (GSVA)^30^ was applied to the gene-level count matrix for three different BRCAness-associated gene sets^31–33^. GSVA enables to assign an enrichment score to each sample and discover distinct BRCAness-like phenotypes from different enrichment patterns. GSVA enrichment scores per sample were computed using the *GSVA* R package with default settings. Gene expression levels were fitted to the recently described network of non-canonical WNT signaling^34^ and network visualization was done with R package *igraph*^35^.

### Identification of master regulator in networks

We pursued an approach called “upstream analysis” which aims to identify master regulators in non-canonical ROR2/WNT11 signaling pathways in MCF-7 breast cancer cells that can be used to find suitable points of intervention for cancer therapy. The upstream analysis strategy comprises three steps: (1) identification of differentially expressed genes (DEGs) based on pairwise comparisons, (2) state-of-the-art promoter analysis to identify relevant transcription factors which are likely to regulate the input DEGs and (3) search for upstream master regulators which are at the very top of the regulatory hierarchy in signal transduction pathways^36^. The main algorithm of the upstream analysis has been described earlier in ^37,38^. In the first step, RSEM estimated counts of samples were processed using the R package DESeq2^39^ to carry out differential expression analysis.

In more detail, DEGs between two conditions (each with triplicate) were identified using the DESeq2 *results* function with the explicit parameters: *alpha* = 0.05, *lfcThreshold = log2(1*.*5)* and *altHypothesis = “greaterAbs”*. Log fold changes were shrunk using the R package *apeglm*^40^ *and genes with adjusted P*-values (Benjamini-Hochberg method) less than 0.05 were considered to be DEGs. In the same step, we also compiled matching sets of non-DEGs for each pairwise comparison, that is, genes whose expression level did not change between tested conditions. The sets of non-DEGs are required for promoter analysis as background sets in the second step of the upstream analysis. For this purpose, we used the *DEseq* function with argument *betaPrior = FALSE* and the *results* function with the explicit parameters: *alpha = 0*.*05, lfcThreshold* = *log2(1*.*5)* and *altHypothesis = “lessAbs”* and genes with adjusted *P*-values less than 0.05 were considered to be non-DEGs.

The upstream analysis was concluded using the ready-to-use workflow called “Upstream analysis (TRANSFAC(R) and TRANSPATH(R))” of the geneXplain platform^38^ web edition 6.2 (https://genexplain.gwdg.de/bioumlweb/) which incorporates the database TRANSFAC(R)^41^ on transcription factors and their DNA binding sites as well as the pathway database TRANSPATH(R)^42^. In our analysis we have run this workflow using our matched lists of DEGs and non-DEGs per comparison as the Inputs for *“Yes gene set*” (target) and *“No gene set”* (background), respectively. We also restricted the search for potential transcription factor binding sites for the promoter analysis to the region of −500 bp upstream (start of promoter) and 100 bp downstream (end of promoter) relative to the transcription start site and left all other parameters at their defaults including an FDR cutoff of 0.05 to retrieve significant master regulators.

### Reverse phase protein array (RPPA)

MCF-7 pcDNA/pROR2 cells were transfected with control (#sc-37007) or WNT11 siRNA (#sc-41120, both santa cruz) as described above. At 48 h post transfection the cells were washed 1x with PBS and lysed in M-PER buffer on ice. Lysates were cleared at 20,000 g, 4° C for 5 min and subjected to RPPA analysis. Briefly, lysates were mixed with 4x printing buffer (125 mM Tris pH 6.8, 10 mM DTT, 4 % SDS, 10 % glycerol), boiled for 5 min at 95°C and printed in triplicates onto nitrocellulose-coated glass slides (Oncyte Avid, Grace Bio-Labs). For blocking, the slides were incubated for 2 h at room temperature in fluorescent western blot blocking buffer (ROCKville) mixed 1:1 with 5 mM NaF, 1mM Na_3_VO_4_ in TBS pH 7.6 followed by incubation with the primary antibodies at 4°C overnight. Antibodies spotted onto RPPA chips were selected for proteins associated with the WNT signaling pathway (**Supplementary Table 1**). Slides incubated without primary antibody served as blank controls. Signals were detected using Alexa Fluor 680 F(ab’)2 fragments of goat anti-mouse or anti-rabbit IgG on an Odyssey infrared imaging system (LI-COR Biosciences) with an excitation wavelength of 685 nm and a resolution of 21 µm. For normalisation, separate slides were stained with Fast Green FCF for total protein quantification. Pre-processing, quality assessment and normalization of RPPA data were performed with the *RPPanalyzer* R package^43^. Differential proteins between the different conditions were identified by fitting linear models and significance calculated using empirical Bayes moderated t-statistics as implemented in the *limma* R package (v3.34.9)^44^.

### Microscopy

Cells were fixed with 4% PFA in PBS (FLUKA) for 15 min and washed twice with PBS. For atomic force microscopy (AFM), cell surface scans were recorded using an MFP-3D atomic force microscope (Asylum Research), equipped with a silicon nitride cantilever (MLCT, *k* = 10 mN·m^-1^, Bruker AFM Probes). Images were recorded in contact mode in PBS at room temperature with a scan rate of 0.3 Hz. Image processing was performed using Gwyddion (http://gwyddion.net/)^45^. For ZO-1 and ROR2 immunofluorescence, samples were blocked in PBS +5% BSA +0.3% Triton X-100 in PBS and stained using an AlexaFluor488-labeled ZO-1 (#339188, Invitrogen) or ROR2 (#FAB20641G, R&D) antibody. Fluorescence was imaged on an upright confocal laser scanning microscope (Olympus, FV 1200) equipped with a water immersion objective (60×, LUMPlanN, NA = 1.0, Olympus). Images were edited in ImageJ. For electron microscopy of cell monolayers, the cells were fixed according to^46^ for 24 h, then embedded in Epon post fixation with 1% OsO_4_, 1.5% uranyl acetate, 1.5% tungstophosphoric acid followed by dehydration with ethanol. A diamond knife was used to prepare ultra-thin sections (50 nm). Micrographs were obtained using a Zeiss EM 912 electron microscope.

### Electric Cell-Substrate Impedance Sensing (ECIS)

ECIS experiments were carried out on a homebuilt setup as described previously^47^. For the measurements an electrode array (8W1E PET ECIS culture ware, Ibidi) was placed in a humidified incubator set at 37 °C with 5% CO_2_. The electrodes were incubated with 200 µL cell culture medium for 1 h before adding 3·10^5^ cells suspended in 300 µL cell culture medium into each well and incubating for 96 h. Impedance was measured at 20 logarithmically spaced frequencies between 10 and 100.000 Hz over time. A cell-electrode model was fitted to the frequency dependent complex impedance spectra from the time point the electrode was covered with a cell layer. The model parameter α, which is inversely proportional to the square root of the cell-substrate distance, was obtained as described previously^48^. Additionally, 1·10^4^ cells per well were seeded onto E-Plates 16 and analysed in the xCELLigence RTCA DP system (Roche) for 48 h in quadruplicates.

### Western blotting, co-immunoprecipitation and mass spectrometry

To analyse protein expression, cells were lysed in RIPA lysis buffer (50 mM Tris, 150 mM NaCl, 0.1 % SDS, 0.5 % sodium deoxycholate, 1 % Triton X-100, pH 7.2) supplemented with protease (Sigma) and phosphatase (Roche) inhibitors, incubated for 15 min on ice, and lysates cleared for 5 min at 20.000 g. Up to 75 µg of protein were separated by SDS-PAGE (8-12% gels), blotted onto nitrocellulose and incubated overnight at 4°C with primary antibodies against WNT11 (#ab31962, abcam), ROR2 (#sc-98486, #sc-80329), RHOA (#sc-418), ROCK1 (#sc-17794), ROCK2 (#sc-398519), PKC (#sc-10800), P-JNK (Thr183/Tyr185, #sc-6254), DAAM1 (#sc-100942), GAPDH (#sc-32233), HSP90 (#sc-13119, all from santa cruz), V5-Tag (#13202), total JNK (#9252, both from cell signaling), TUBA (#05-829, Millipore). Membranes were incubated with HRP-coupled secondary antibodies (santa cruz, cell signaling) for 1 h at RT and chemiluminescence detected with ECL Prime (GE Healthcare) at the LAS-1000 (Fujifilm), Amersham Imager 600 (GE Healthcare) or ChemoStar Touch Imager (Intas). Image J software was used for densitometric quantification. For co-immunoprecipitation, cells were transfected with V5-tagged WNT11 and 24 h post transfection proteins were crosslinked for 30 min with 1 mM DSS (Thermo Fisher) in 1 ml PBS + 1mM MgCl_2_. Cells were washed once in PBS and lyzed for 30 min on ice in 50 mM Tris/HCl pH 8.0, 150 mM NaCl, 1% NP-40 supplemented with protease inhibitors (Sigma). Lysates were cleared for 10 min at 16,000 g and 500 µg protein were incubated with 1 µg normal rabbit IgG (#2729) or V5 antibody (#13202, both cell signaling) for 16 h at 4°C. Antibody-protein complexes were incubated for 2 h at 4°C with protein A/G agarose beads (#sc-2003, santa cruz) and spun down at 1,700 g for 1 min. Signals were visualized by western blot as described above using the confirmation-specific mouse anti-rabbit IgG (#3678, cell signaling) for V5 detection.

### Gene expression analysis

Total RNA was extracted with the High Pure RNA isolation kit (Roche) and 1 µg reversely transcribed into cDNA (iScript cDNA synthesis kit, Bio-Rad). Gene expression was assessed from 10 ng cDNA at the ABI 7900 HT system using SYBR green detection and the SDS (v2.4) software (Applied Biosystems). For normalization the two housekeeping genes *HPRT1* and *GNB2L1* were used. Primer sequences are given in **Supplementary Table 2**.

### Flow cytometry

Cells were washed once in PBS, blocked for 15 min in PBS +1% FCS and stained for 20 min with either a ROR2 Alexa Fluor 488-conjugated antibody (#FAB20641G) or the respective isotype control (#IC003G, both R&D). Stained cells were measured on an Attune NxT flow cytometer (ThermoFisher) and the data were analyzed with FlowJo (v10.6.1).

### Cell invasion and migration

Cell invasion was measured from 1·10^5^ cells per well in triplicates in a modified Boyden chamber and related to the unstimulated control^49^. Cells were either pre-incubated for two hours with IWP-2 (Cayman Chemical), WNT-C59 (Tocris), sFRP1 (R&D systems), DKK1 (R&D systems), JNK inhibitor (SP600125, Tocris), RHO inhibitor Rhosin (Tocris), or pre-treated for 24 h with the indicated siRNAs.

### RHOA activity assay

MCF-7 cells were transfected with the indicated siRNA as described above and 48 h post transfection incubated in RPMI-1640 +1% FCS overnight. The next day, samples were stimulated with RPMI-1640 +10% FCS for 30 min. Lysate preparation and measurement of RHOA-GTP levels were carried out with the RHOA G-LISA activation assay kit (#BK124, Cytoskeleton) according to the manufacturer’s instructions.

### Statistics

Results are displayed as means ± standard deviation (SD). Statistical analyses were carried out with GraphPad Prism (v8.4.2) using a two-sided paired/unpaired *t*-test unless stated otherwise. *p* < 0.05 was considered statistically significant. All experiments were carried out as independent biological replicates as indicated by the “n” number in the Figure legends. All figures in the current study were either created with GraphPad Prism (v8.4.2) or generated in R (v3.6.2). The *ggsurvplot* function from the *survminer* R package was used to generate Kaplan–Meier survival plots. For heatmap visualizations, the functions *heatmap*.*2* and *pheatmap* from the R packages *gplots* (v3.1.0, https://CRAN.R-project.org/package=gplots) and *pheatmap* (v1.0.12, http://CRAN.Rproject.org/package=pheatmap) were used, respectively.

## RESULTS

### High ROR2 expression is associated with early metastasis in breast cancer patients

Based on the observations of active non-canonical WNT signaling in breast cancer tissue, we compared the expression levels of the four non-canonical WNT co-receptors *ROR1, ROR2, PTK7* and *RYK* in normal and cancerous breast tissue using the IST Online™ and TNMplot database (**Fig. 1a, Supplementary Fig. 1**). While *ROR2* as well as *PTK7* were expressed at higher levels in cancerous tissue, *RYK* expression seemed to be downregulated, and no clear trend was observed for *ROR1*. We then investigated whether the expression levels of the receptors were correlated with the clinical prognosis using a compendium dataset which comprises gene expression data of primary breast cancers from ten public datasets with a total of 2,075 patients^12^. Indeed, the expression levels of *ROR1, ROR2, PTK7* and *RYK* were negatively associated with metastasis-free survival (MFS) of the patients, although the effect was not statistically significant for the latter (**Fig. 1b**). These observations suggested that *PTK7, ROR1* and *ROR2* contribute to an aggressive cancer cell phenotype that promotes early metastasis formation. Since the strongest effect was seen for *ROR2*, we focussed our further analyses on this receptor.

**Figure 1:**
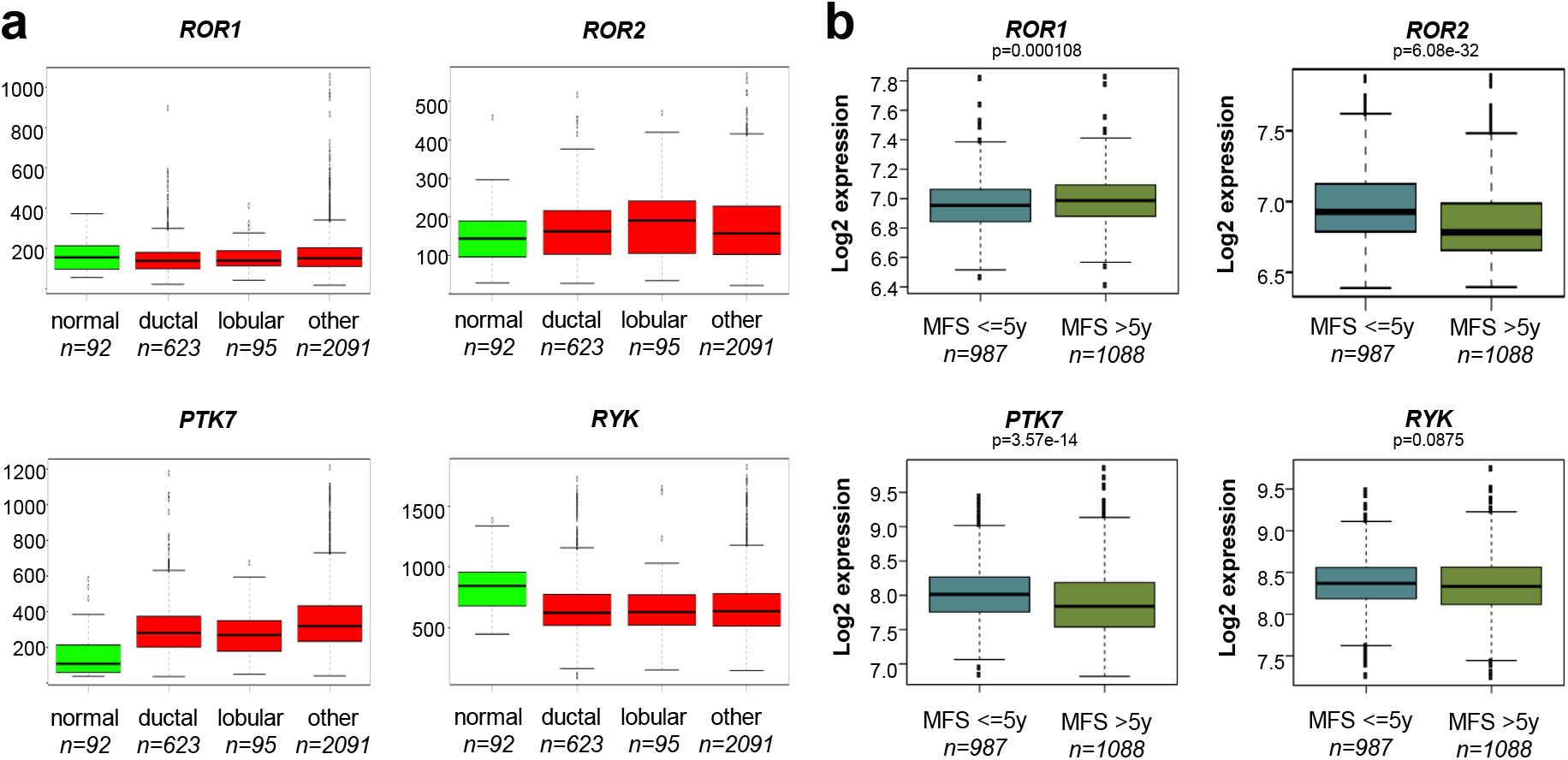
Elevated levels of ROR2 and PTK7 in primary breast cancer are associated with early metastatic spread. ***a***, Gene expression of the four non-canonical WNT co-receptors *ROR1, ROR2, PTK7* and *RYK* in normal breast (green) or breast cancer (red) from the IST Online™ database (ist.medisapiens.com). ***b***, Microarray gene expression data from 2,075 breast cancer primary tumors were correlated with either poor (≤ 5 years) or better (> 5 years) metastasis-free survival (MFS). Significance was calculated with a t-test. Boxes represent the 25-75^th^ percentiles with the line at the median. Outliers are marked as small dots.

### ROR2 expression induces an aggressive cancer cell phenotype linked to BRCAness

Next, we aimed to unravel the molecular events that underlie the unfavorable prognostic role of *ROR2*. Since ROR1 and ROR2 are known to interact and compensate for each other, we choose to limit potential cross-reactivity by stably overexpressing ROR2 in the ROR-negative^11^ human breast cancer cell line MCF-7 (**Fig. 2a**). Interestingly, overexpression of ROR2 changed the molecular characteristics of the cell line from the rather benign luminal A to the highly-aggressive basal-like subtype as calculated from the PAM50 gene signature (**Fig. 2b**). This is in correspondence with their increased invasive and migratory potential as well as with observations made in mouse models^11,14,16^. Since some triple-negative breast cancers share the high-grade genomic instability observed in BRCA1/2-mutant cancers^50^, the so-called BRCAness, we were interested in whether ROR2 could be linked to these changes. Indeed, the gene expression profile of the ROR2-overexpressing cells showed a higher enrichment for three independently published BRCAness signatures^31–33^ (**Fig. 2c**). Moreover, the cells showed increased susceptibility to treatment with the poly (ADP-ribose) polymerase (PARP) inhibitor olaparib (**Fig. 2d**), underlining that ROR2 induces a BRCAness-like phenotype in breast cancer cells.

**Figure 2:**
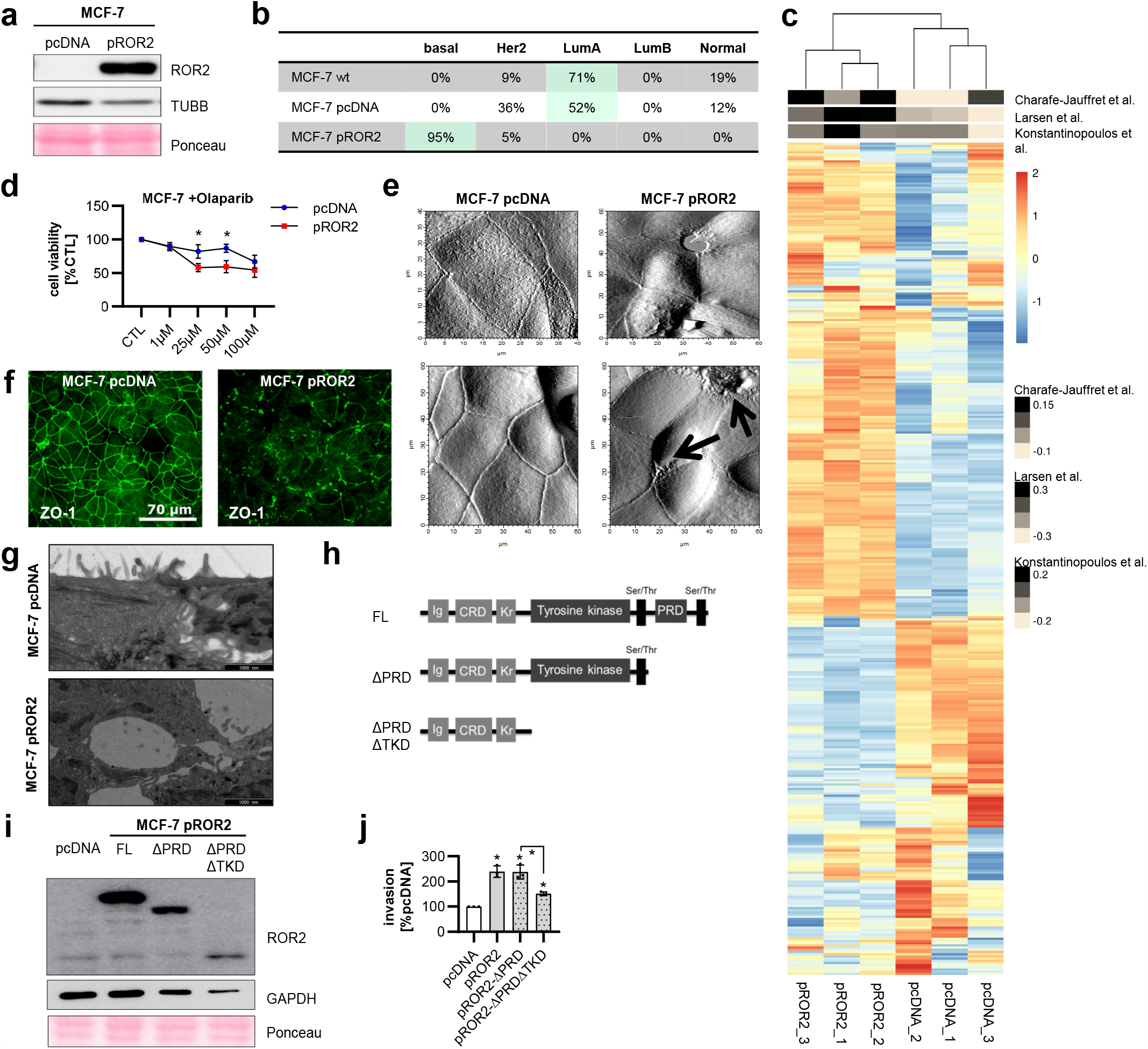
Expression of ROR2 induces a highly invasive phenotype in breast cancer cells. ***a***, MCF-7 human breast cancer cells were transfected either with an empty vector (pcDNA) or with a hROR2 overexpression plasmid (pROR2). Transfection efficiency was confirmed by western blot. ***b***, Determination of the molecular breast cancer subtype based on RNA-Seq data from the indicated cell lines using the PAM50 gene signature. ***c***, GSVA analysis for three independently published BRCAness gene expression signatures. ***d***, MTT assay: Cells were treated for 96 h with the indicated concentrations of olaparib (mean±SD, n=3, *p<0.05). ***e***, Atomic force microscopy (AFM) of pcDNA and pROR2 cells. ***f***, Immunofluorescence staining of the tight junction marker ZO-1. ***g***, Electron microscopy of tight junctions (scale bar: 1 µm). ***h***, Schematic overview of the pROR2 C-terminal deletion constructs. ***i+j***, C-terminal deletion constructs were overexpressed in MCF-7. Expression was confirmed by western blot *(i)* and the invasion rate measured in Boyden chambers *(j)* (mean±SD, n=3, *p<0.01).

In line with the rather mesenchymal, highly motile phenotype typically observed in basal-like cancer cells, ROR2 overexpression resulted in apparent defects in cell-cell-contacts with large gaps in confluent cell layers and membrane ruffles at cellular junctions (**Fig. 2e**). These morphological changes were not caused by alterations in the actin cytoskeleton, which seemed comparable in both cell types (**Supplementary Fig. 2**). In contrast, the distribution of the scaffolding protein ZO-1, which is required for the assembly of tight junctions, was severely disrupted. While ZO-1 was equally distributed at the plasma membrane in control cells, it clustered in large dots in pROR2 cells (**Fig. 2f**). A closer imaging of the cells by electron microscopy confirmed severe structural defects in cellular tight junctions which were almost completely absent (**Fig. 2g**).

In its C-terminus ROR2 harbors an active tyrosine kinase domain (TKD)^51^ as well as a proline-rich domain (PRD) that potentially mediate protein-protein interactions. In order to gain insight into the molecular mechanisms behind the aggressive function of ROR2, we transfected MCF-7 cells with *ROR2* constructs which either lacked the PRD (ΔPRD), or both PRD and TKD (ΔPRDΔTKD) (**Fig. 2h+i, Supplementary Fig. 3a+b**). Using Boyden chamber assays we observed that only the double mutant showed a strong reduction in its invasive potential (**Fig. 2j**), suggesting that the tyrosine kinase function of ROR2 is essential for its invasion-promoting function.

**Figure 3:**
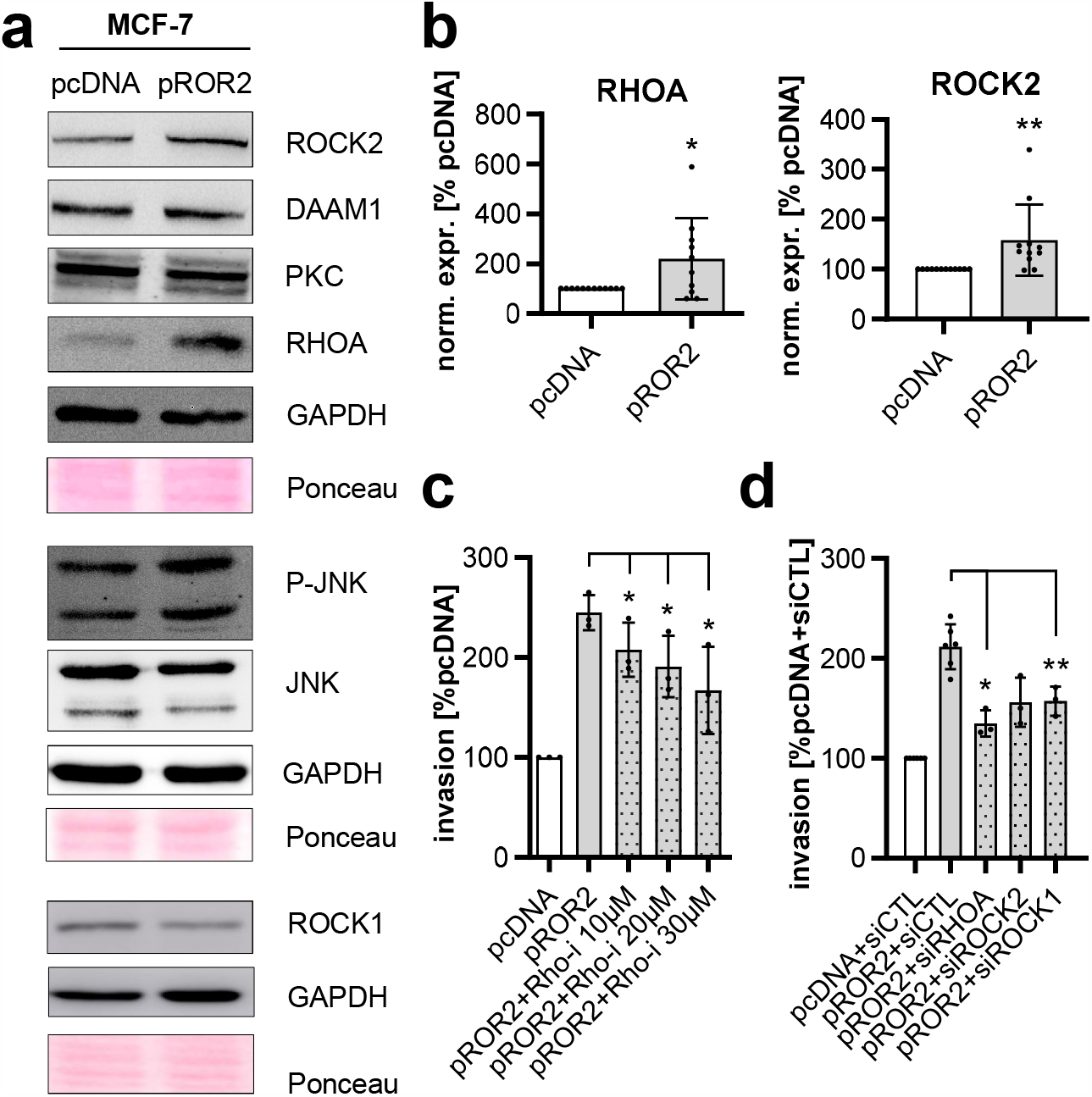
RHOA/ROCK mediate ROR2-induced tumor invasion. ***a***, pcDNA and pROR2 cells were characterized for the expression of non-canonical WNT signaling proteins by western blot. ***b***, Densitometric quantification of RHOA and ROCK2 in MCF-7 pROR2 normalized on GAPDH expression (mean±SD, n=10-11, *p<0.05, **p<0.01). ***c***, Invasion assay: MCF-7 pROR2 cells were treated for 96 h with the indicated concentration of Rhosin, a RHO inhibitor (mean±SD, n=3, *p<0.05). ***d***, MCF-7 pROR2 cells were transfected with the indicated siRNAs (10 nM) and cell invasion was measured in Boyden chambers (mean±SD, n=3, *p<0.05, **p<0.01).

### ROR2 enhances tumor invasion via RHO/ROCK signaling

We had previously shown that overexpression of ROR2 does not affect the levels or localization of β-catenin in breast cancer cells and thus does not activate canonical WNT signaling^11^. To identify the oncogenic signaling responsible for the tumorigenic functions of ROR2, we characterized the ROR2-overexpressing cells by western blot for the expression of common non-canonical WNT signaling molecules (**Fig. 3a**). The most robust effect was a significant upregulation of the small GTPase Ras homolog family member A (RHOA) and its effector RHO-associated coiled-coil containing protein kinase 2 (ROCK2) in pROR2 cells (**Fig. 3b**). Similarly, there was a trend to increased phosphorylation of the c-Jun N-terminal kinase (P-JNK) consistent with our own previous observations^11^. Taken together, these results suggested that ROR2 activates WNT/PCP signaling.

To investigate whether the increased expression of RHOA is involved in the invasion-promoting role of ROR2, we treated MCF-7 pROR2 cells with the RHO inhibitor Rhosin which blocks the activity of RHOA by inhibiting its interaction with its guanine nucleotide exchange factors (GEFs). As expected, this reduced the pro-invasive function of ROR2 in Boyden chamber assays in a concentration-dependent manner (**Fig. 3c**). Likewise, the same effect was seen after specific knockdown of RHOA, ROCK1 and ROCK2 by siRNA (**Fig. 3d**). This suggested that even though the expression of ROCK1 had initially not been found to differ between cells transfected with empty vector or pROR2, the ROR2-induced increase in cancer cell invasiveness involved both ROCK1 and ROCK2.

### ROR2 triggers expression of its putative non-canonical WNT ligands

Next, we searched for potential non-canonical WNT ligands that could activate ROR2-induced WNT/PCP signaling. A comparison of MCF-7 empty vector and ROR2-overexpressing cells by RNA-Seq identified 2,860 differentially expressed genes (DEGs) in both cell lines^12^. To narrow down our search, we specifically filtered that list for genes that are part of the recently constructed network representing non-canonical WNT signaling^34^. Our analyses revealed the non-canonical WNT ligand *WNT11* as one of the most highly upregulated genes in the ROR2-overexpressing cells (**Fig. 4a**). The *WNT11* upregulation was confirmed at the protein level and was independent of WNT5A stimulation, the established ligand of ROR2 (**Fig. 4b**). An analysis of the expression of other non-canonical WNT genes in MCF-7 cells revealed that there was no upregulation of any other typical non-canonical WNT ligand including *WNT4, WNT5A* or *WNT6* (**Fig. 4c**). A similar increase in *WNT11* levels was confirmed for BT-474 cells overexpressing ROR2 (**Fig. 4d**). The same trend was also observed in MDA-MB-231 cells, although it did not reach statistical significance. Other breast cancer cells reacted with the induction of different non-canonical WNT ligands, such as *WNT5A* in SK-BR-3 and *WNT6* in T-47D cells. Together, this supports the assumption that ROR2, when overexpressed, upregulates its own potential ligands in a cell context-dependent manner.

**Figure 4:**
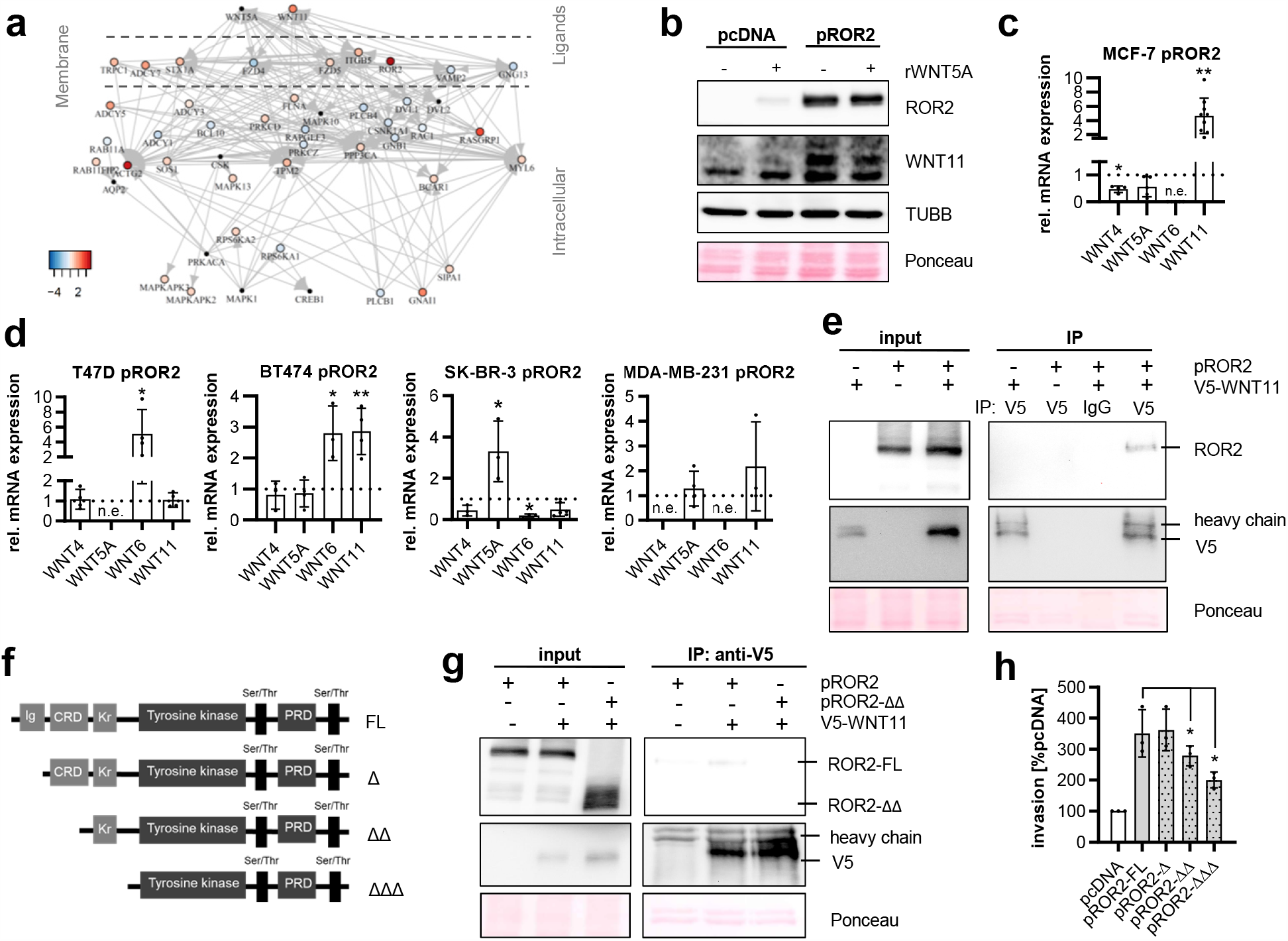
WNT11 is a novel ligand for ROR2 in humans. ***a***, RNA-Seq of MCF-7 pROR2 cells: Network of differentially expressed genes associated with non-canonical WNT signaling grouped according to their cellular localization. ***b***, MCF-7 cells were stimulated for 24 h with rWNT5A (100ng/ml) and WNT11 expression was analyzed by western Blot. ***c+d***, Expression of the non-canonical WNT ligands was measured by qRT-PCR in MCF-7 *(c)* or the indicated ROR2-overexpressing human breast cancer cell lines *(d)* (mean±SD, n=3-9, *p<0.05, p<0.01, n.e.= not expressed). ***e***, Co-immunoprecipitation (Co-IP) of V5-WNT11 in MCF-7 pROR2 cells detects ROR2 by western blot. ***f***, Schematic representation of the ROR2 N-terminal deletion constructs. ***g***, Co-IP of V5-Wnt11 in MCF-7 expressing either pROR2-FL or pROR2-ΔΔ. ***h***, Cell invasion assays of MCF-7 expressing N-terminal ROR2 deletion constructs (mean±SD, n=3, *p<0.0001).

### WNT11 is a novel ligand for ROR2 in humans

Based on these results, we hypothesized that WNT11 might act as a novel ligand for ROR2. WNT11 has already been identified as a ligand for ROR2 in xenopus gastrulation^52^. However, it remained unknown whether WNT11 can also interact with ROR2 in humans and whether it might be responsible for its tumor-supporting function. To demonstrate that WNT11 indeed binds to human ROR2, we transiently transfected MCF-7 ROR2-overexpressing cells with a functionally active V5-tagged WNT11 (**Supplementary Fig. 3c**). ROR2 was co-immunoprecipitated with V5-WNT11 (**Fig. 4e**), thus confirming their interaction.

ROR2 harbors three extracellular domains: an immunoglobulin-like (Ig-like), a cysteine-rich as well as a Kringle domain. In Xenopus it has been shown that WNT proteins preferentially interact with the extracellular region of ROR2, in particular the CRD^53^, while the Kringle domain seems to be important for receptor heterodimerization^54^. To investigate which domain is required for the pro-invasive function of ROR2, we cloned serial deletion constructs of ROR2 lacking the three extracellular domains and overexpressed them in MCF-7 cells (**Fig. 4f**). The deletion did not interfere with the localization of ROR2 which was consistently expressed at the plasma membrane in all transfected cells (**Supplementary Fig. 3d**). Deletion of the first two domains abolished the interaction of V5-WNT11 and ROR2 in co-immunoprecipitations (**Fig. 4g**). While the lack of the first, Ig-like extracellular domain, did not affect the invasion-promoting effect of ROR2, a significant reduction in cancer cell invasiveness was observed when overexpressing ROR2 lacking the CRD (**Fig. 4h**). This trend was even more pronounced upon deletion of the Kringle domain. In summary, these observations indicated that WNT11 is a novel ligand for ROR2 interacting with its CRD and that signal transduction involves receptor heterodimerization.

### WNT11 is responsible for the tumorigenic functions of ROR2 in breast cancer cells

We then aimed at further elucidating the consequences of the WNT11/ROR2 interaction for tumor cell function. Treatment of ROR2-overexpressing cells with soluble WNT ligand antagonists such as sFRP1 or DKK1 as well as an inhibitor blocking non-canonical WNT-JNK signaling significantly inhibited ROR2-induced cancer cell invasiveness (**Fig. 5a**). In line with this, the two porcupine inhibitors IWP-2 and WNT-C59, which block the secretion of WNT ligands, had the same effect (**Fig. 5b**), underlining that a cell-intrinsic WNT ligand seemed to be responsible for the pro-invasive effect of ROR2. To confirm the involvement of WNT11 in this process, we generated MCF-7 pROR2 cells with a stable, shRNA-mediated knockdown of WNT11 (**Supplementary Fig. 4a**). The invasiveness of these cells was significantly impaired (**Fig. 5c**). Similarly, a transient reduction of WNT11 expression by siRNA (**Supplementary Fig. 4b**) resulted in decreased tumor cell invasion, an effect that was rescued by addition of recombinant WNT11 (**Fig. 5d**), thus confirming that WNT11 mediates a major part of the invasion-promoting effect of ROR2.

**Figure 5:**
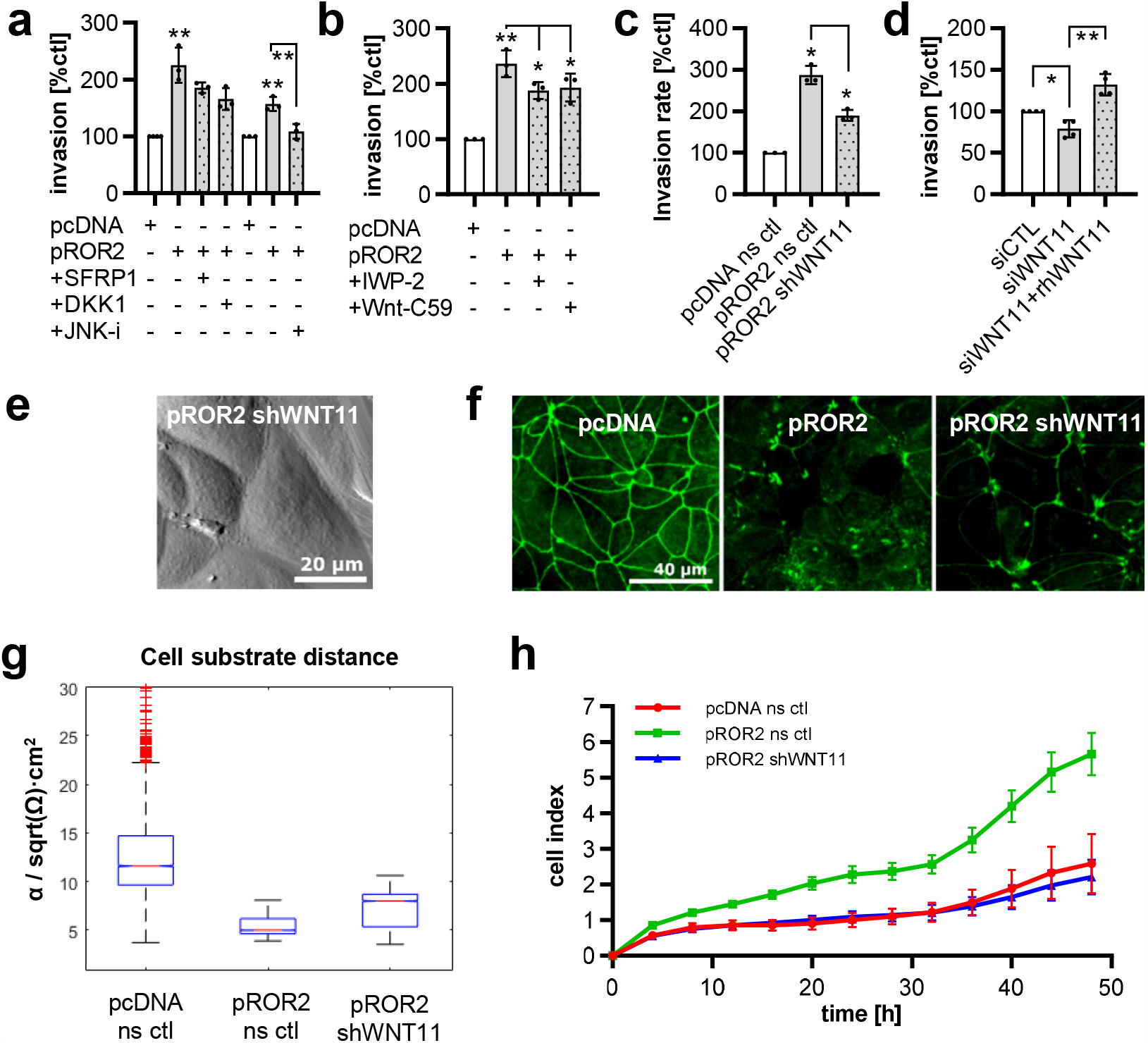
WNT11 mediates the pro-tumoral effects of ROR2. ***a+b***, Invasion assay: MCF-7 pcDNA or pROR2 cells were treated with WNT inhibitors *(a)* or Porcupine inhibitors *(b)* (mean±SD, n=3, *p<0.05, **p<0.01). ***c***, Invasion assay of MCF-7 pROR2 cells stably expressing a non-sense control (ns ctl) or WNT11 (shWNT11) shRNA (mean±SD, n=3, *p<0.01). ***d***, Invasion of MCF-7 pROR2 cells transfected with control (siCTL) or WNT11 siRNA (siWNT11) +/- rhWNT11 (100 ng/ml) was assessed in Boyden chambers (mean±SD, n=3, *p<0.05, **p<0.001). ***e***, AFM of MCF-7 pROR2 shWNT11 cells. ***f***, Immunofluorescence for the tight junction protein ZO-1. ***g***, ECIS measurements (box: 25-75^th^ percentile, line at median). ***h***, xCELLigence measurements (mean±SD). Shown is one representative example out of three independent experiments.

When the growth pattern and shape of MCF-7 pROR2 cells with stable WNT11 knockdown were analyzed, the cells resembled the empty vector control cells (see **Fig. 2c**) with only few membrane ruffles or gaps in the confluent cell layer (**Fig. 5e**). This amelioration of the defects in cell-cell-contacts was also mirrored in the distribution of ZO-1, which started to line again the cell-cell borders upon WNT11 depletion comparable to the empty vector control cells (**Fig. 5f**). To evaluate the significance of these changes, we performed Electric Cell-Substrate Impedance Sensing (ECIS) measurements that permit the evaluation of the behavior of adherent cells, and are influenced by cellular morphology, adhesion and proliferation. Using this method, we were able to demonstrate that the aggressive phenotype of the pROR2 WNT11 knockdown cells showed a reversion towards the control cells, including a significantly higher cell substrate distance as well as a lower cell index compared to the ROR2-overexpressing cells (**Fig. 5g+h**).

### WNT11 mediates ROR2 signaling

Since we had identified RHOA and ROCK2 as critical signaling mediators for ROR2-induced invasion, we were interested in whether their activation was dependent on WNT11. Indeed, siRNA-mediated knockdown of WNT11 in ROR2-overexpressing cells antagonized the observed increase in RHOA, although no significant effect on ROCK2 expression was detectable (**Fig. 6a**). Using an ELISA for the active, GTP-bound form of RHOA, we showed that this regulation was not only present at the level of total RHOA protein, but that ROR2 triggered the activation of RHOA, an effect that was effectively counteracted by knockdown of WNT11 (**Fig. 6b**).

**Figure 6:**
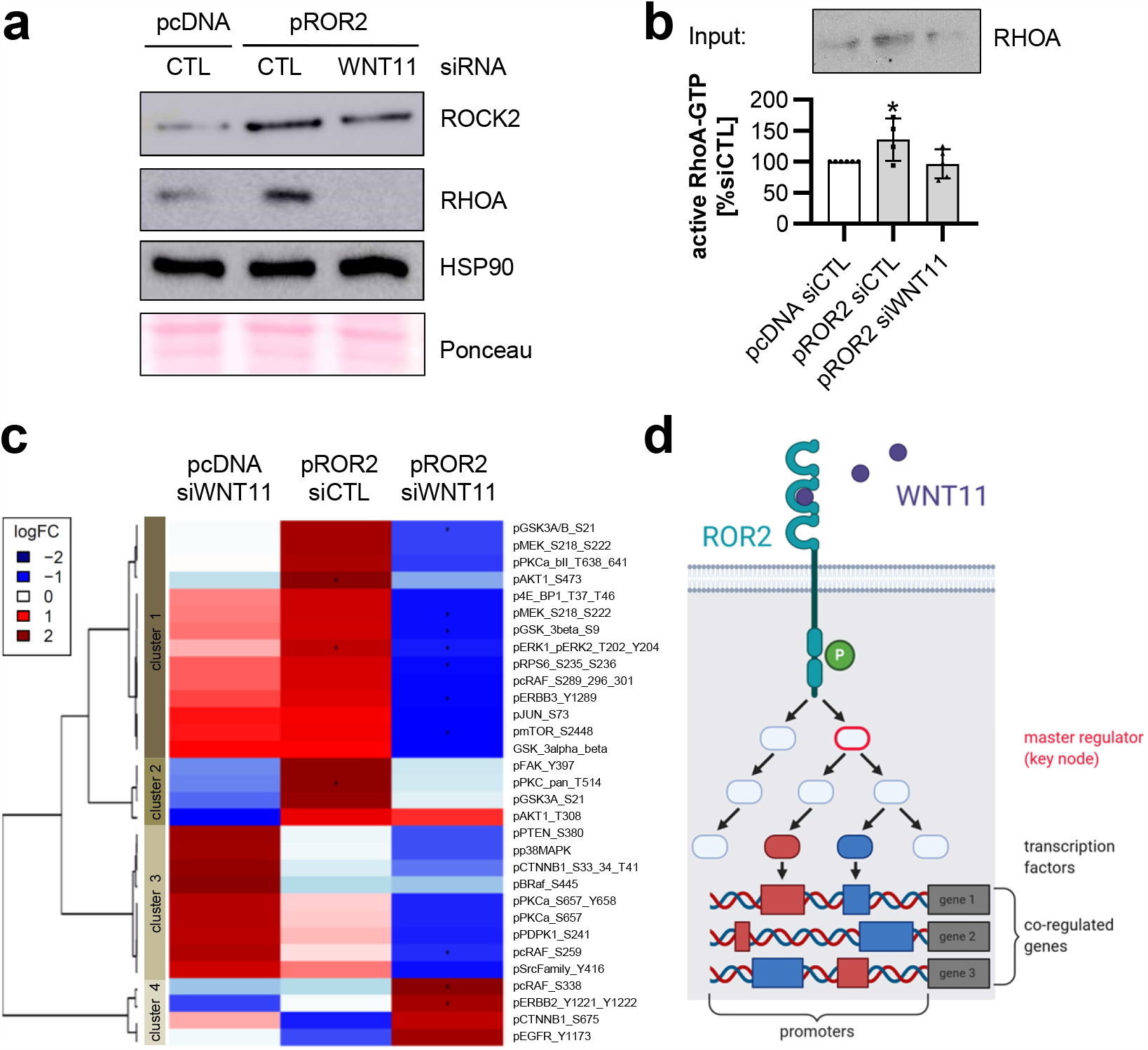
WNT11 mediates ROR2 signaling. ***a***, Western blot: RHOA and ROCK2 in MCF-7 pROR2 cells treated with control siRNA (siCTL) or siRNA against WNT11 (siWNT11). ***b***, Levels of active RHOA-GTP in the indicated cells were assessed by ELISA (mean±SD, n=5, *p<0.05). ***c***, MCF-7 pcDNA and pROR2 siCTL/siWNT11 cells were characterized by RPPA for phospho-proteins associated with the WNT signaling pathway (n=3, *p<0.05). ***d***, Schematic representation of the master regulator analysis of ROR2/WNT11 signaling. The illustration was created with BioRender.com.

To gain further insight into additional pathways regulated by ROR2/WNT11, we characterized ROR2-overexpressing cells with siRNA-mediated WNT11 knockdown by RNA-Seq as well as reverse phase protein array (RPPA). The RPPA chip contained antibodies against phospho-proteins involved in WNT signaling and associated pathways. Expression levels are presented in a heatmap in **Fig. 6c**. The array results showed that in particular proteins in cluster 1 and 2 were highly upregulated in pROR2 cells compared to empty vector control cells, and that these expression changes were reverted upon knockdown of WNT11, indicating that these sets of proteins were regulated through ROR2/WNT11.

We then aimed at identifying common master regulators that could mediate the activation of sets of these ROR2/WNT11 targets (**Fig. 6d**). In order to do so, we first identified differentially expressed genes (DEGs) between empty vector control and pROR2 cells based on the RNA-Seq results. Next, the promoters of DEGs were analyzed for enriched transcription factor binding sites to define sets of relevant transcription factors and relate them to upstream signal transduction pathways based on the regulatory pathway database TRANSPATH. The search for upstream signaling molecules that are responsible for regulating these sets of transcription factors finally identified the responsible master regulators. The master regulator analysis of empty vector and pROR2 cells revealed *PIK3CA and RHOA* as master regulators of signaling in ROR2-overexpressing cells (**Supplementary Table 3, Supplementary Fig. 5**). This fits to the results of the RPPA characterization with several signaling molecules of the PI3K pathway present in the WNT11-regulated clusters 1 and 2 (e.g. P-Akt, P-mTOR, P-RPS6) (**Fig. 6c**). Moreover, it suggests that WNT11 is able to activate PI3K signaling, indicating that it might be responsible for the activation of PI3K in pROR2 cells. In line with this hypothesis, *PIK3CA* was no longer detectable as a master regulator in pROR2 cells after knockdown of WNT11 (**Supplementary Table 4**). Taken together, these results imply that WNT11 is able to activate tumor-promoting signaling pathways in breast cancer cells via its interaction with ROR2.

### WNT11/ROR2 are highly expressed in breast cancer brain metastases and are associated with poor patient survival

While studies have suggested hyperactive WNT signaling in primary breast cancers, it is still not clear whether the same holds true for metastases which are known to differ from the primary^9^. We therefore collected samples from 31 patients with brain metastases and characterized them by RNA-Seq. With the obtained data, we performed a pathway enrichment analysis looking at three different gene sets, either associated with canonical, non-canonical, or regulation of WNT signaling^34^ (**Fig. 7a**). Interestingly, genes associated with non-canonical WNT signaling were highly enriched in almost all samples, and the enrichment was more pronounced than for canonical WNT signaling, the cluster of patients with the highest enrichment was characterized by significantly shorter overall survival.

**Figure 7:**
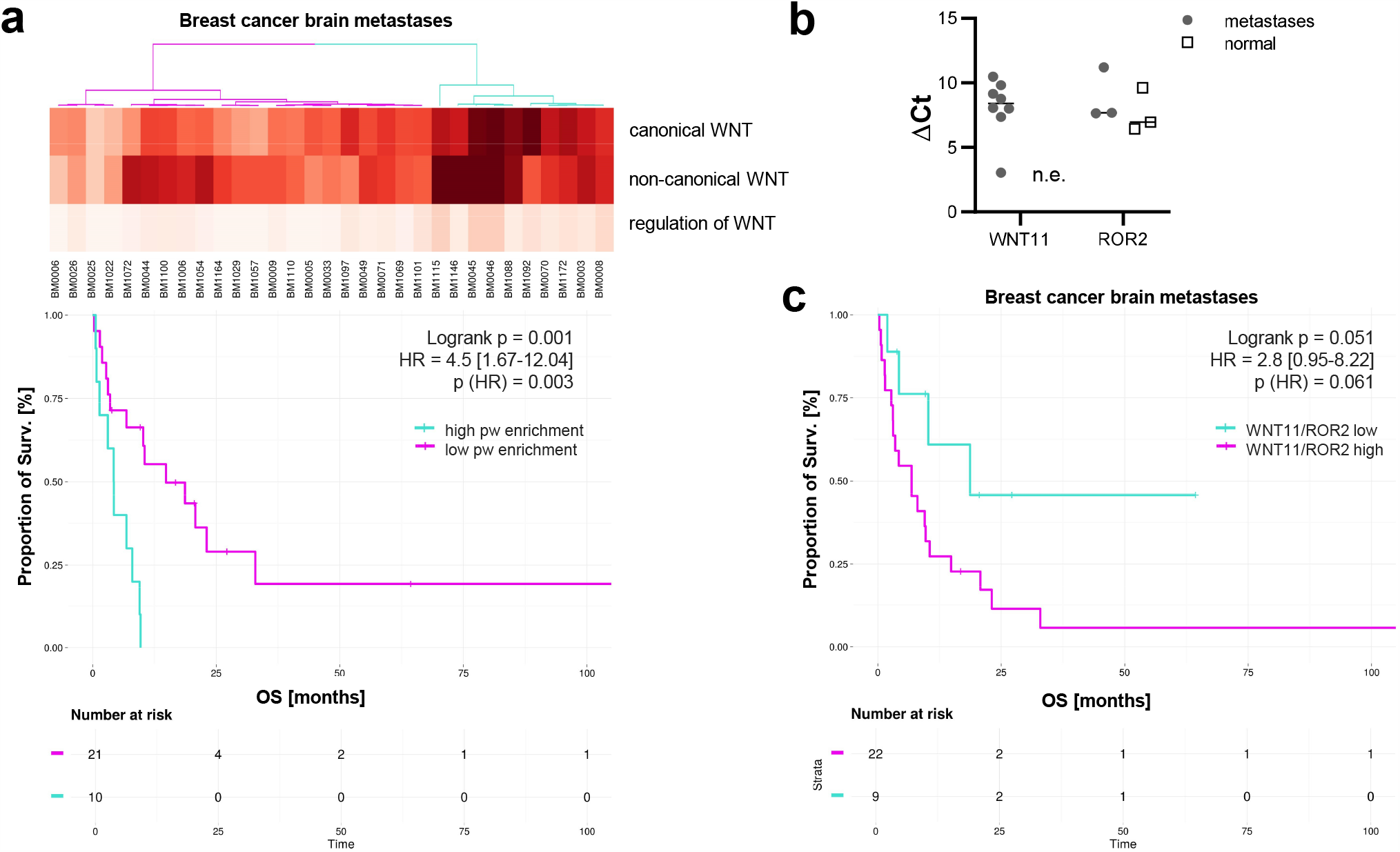
ROR2/WNT11 are expressed in metastatic breast cancer and associated with poor survival. *a*, Pathway enrichment for RNA-Seq data from 31 patients with brain metastases given for different WNT subpathways. Significance was calculated with a log rank test. ***b***, qRT-PCR: Expression of *ROR2* and *WNT11* in samples of human brain metastases (line at median, n.e.= not expressed). ***c***, Kaplan-Meier survival curves showing the OS of metastatic patients based on their averaged *WNT11* and *ROR2* expression. The separation high/low was computed based on an optimal cutoff using the maxstat method^55^. Significance was calculated with a log rank test.

Next, we isolated RNA from seven metastases as well as normal brain tissue and analyzed the expression of *ROR2* and its ligand *WNT11* by quantitative real-time PCR (**Fig. 7b**). While *ROR2* was present in half of the metastases and control samples, *WNT11* was undetectable in normal brain tissue, but highly expressed in all metastases, thus further pointing towards its important role in metastatic growth. To confirm the role of *ROR2/WNT11* in metastasis, we correlated the expression levels of *ROR2* and *WNT11* in the metastatic tissue with patient outcome using the gene expression data obtained by RNA-Seq. Indeed, patients with high expression of *ROR2* plus *WNT11* had the shortest overall survival (**Fig. 7c**). Although the results did not quite reach significance due to the limited number of patient samples, the results do indicate that WNT11, as a novel ligand for ROR2, also exerts an unfavorable role in brain metastasis of breast cancer patients *in vivo*.

## DISCUSSION

Previously, we had described a gene expression signature comprising 76 genes regulated by the WNT co-receptor ROR2 which grouped primary breast cancer patients into two clusters with significant differences in MFS^12^. Moreover, we had detected high expression levels of ROR2 in breast cancer brain metastases^10^, suggesting its active involvement in tumor progression. Now, we explored the molecular mechanisms underlying these observations and showed that ROR2 confers an aggressive phenotype to breast cancer cells that is linked to basal-like features and a high invasive potential. We identified WNT11 as a novel ligand for ROR2 that activates WNT/PCP signaling and is responsible for the induction of tumor invasion. WNT11 is highly expressed in brain metastases and is associated with poor patient survival when co-expressed with ROR2. ROR2-positive tumor cells are characterized by a BRCAness-like gene expression signature, and an enhanced susceptibility to PARP inhibition, thereby opening new possibilities for targeted treatment.

ROR2 has been reported to act either as a tumor suppressor or an oncogene depending on the type of tumor^56^. This is not surprising since WNT signaling responses are highly cell context-dependent and involve a multitude of WNT subnetworks that result in diverse functional outcomes. WNT ligands have been shown to compete for receptor binding on the cell surface^57^. Hence, the combination of available ligand, receptor and co-receptor seem to dictate which subnetworks are activated^3,4,57^. Our own results indicate that in breast cancer especially non-canonical WNT signaling is highly active 10^10,11,^12. Correspondingly, a negative correlation between ROR2 and active canonical WNT signaling was revealed in human breast tumors based on the TCGA database^16^. So far, available evidence indicates that ROR2 acts as an oncogene in breast cancer. *In vitro* studies have revealed a stimulatory function of ROR2 on breast cancer cell invasion and migration, mostly through induction of an EMT-like cancer cell phenotype^11,14,15^. Moreover, the treatment of MCF-7 cells with increasing doses of tamoxifen generated resistant clones that exhibited an EMT-like phenotype and were characterized by elevated ROR2 levels^58^. This fits our detection of morphological alterations with a switch to the aggressive, highly motile basal-like subtype in ROR2-overexpressing cells. In mice, ROR2-positive tumors are associated with accelerated tumor growth and a shorter survival time, confirming the tumor-promoting role of ROR2 *in vivo*^16,59^. Interestingly, overexpression of ROR2 induced a BRCAness-like gene expression profile in breast cancer cells. To our knowledge, ROR2 has not yet been associated with genomic instability, thus necessitating further research to understand this finding. However, the observation that ROR2-overexpressing cells become susceptible to PARP inhibition paves the way for a new approach for treating ROR2-positive tumors.

Although initial studies in Xenopus and C. elegans had identified WNT binding domains in ROR2^53,60^, and a physical interaction of ROR2 with WNT11 had been described in Xenopus^52^, WNT5A was still believed to be the sole ligand for ROR2 in humans^56^. The newly identified interaction between human ROR2 and WNT11 suggests a similar promiscuity for ROR2 in ligand binding compared to the FZD receptors. The observation that ROR2 is able to upregulate the expression of several non-canonical WNT ligands in a cell line-specific context, highlights WNT6 as another potential ligand for ROR2. In contrast, WNT4 was not differentially expressed upon ROR2 overexpression in the investigated breast cancer cell lines. However, it cannot be excluded that it might act as a ROR2 ligand in a particular context.

*It remains unclear whether WNT11 stimulation couples ROR2 to additional co-receptors (e*.*g*. FZDs) as it has been observed for WNT5A^57^. Interestingly, while the deletion of the Ig and CRD domain of ROR2 decreased its invasion-promoting effect, an additional reduction was observed upon deletion of the Kringle domain. As the Kringle domain has been shown to contain a lysine-binding site that could potentially mediate receptor-receptor interactions^61^, this observation might point to the involvement of other receptors in the WNT11/ROR2 complex. ROR2 is able to heterodimerize with PTK7 or VANGL2 to stimulate vertebrate WNT/PCP signaling^17,18^, raising the question whether apart from the FZDs both might also serve as candidates for members of a possible complex formation.

*WNT11* was reported to be highly amplified in 6.2% of breast cancer patients^13^ and observed in specific tumor cell subpopulations^16^. Moreover, *WNT11* was recently identified as one of three WNT genes mutated specifically in breast cancer metastases compared with matched primary tumors^9^. While our data reveal a high expression of *WNT11* in brain metastases and poor patient survival when co-expressed with ROR2, the effect of WNT11 signaling might vary in different receptor contexts. In our cell line model WNT11/ROR2 signaling resulted in activation of RHOA/ROCK and thus elicited a typical WNT/PCP response. This is in line with studies of WNT11 signaling in Xenopus and mouse^16,52^. The observation was further confirmed by our network analysis which identified RHOA as well as PIK3CA as master regulators of ROR2 signaling. Since PIK3CA was lost as a master regulator upon WNT11 knockdown, this suggests that WNT11 might integrate WNT and PI3K/AKT signaling. Although this requires further proof, similar observations have been claimed for prostate cancer^62^.

Taken together, we show for the first time that ROR2 can trigger an autocrine, invasion-promoting signaling response via RHO/ROCK by inducing expression of its own ligand, WNT11, in human breast cancer. Obviously, the notion of WNT5A as the sole ligand for human ROR2 no longer holds true which opens novel insights in the regulation of non-canonical WNT signaling in cancer. ROR2/WNT11 upregulation is especially relevant in the metastatic situation since it confers poor prognosis on the one hand, but also sensitivity towards PARP inhibitors on the other, thus providing a promising therapeutic target worth further exploration.

## ACKNOWLEDGEMENTS

This work was supported by the Deutsche Forschungsgemeinschaft (DFG, German Research Foundation - project 424252458), the Göttinger Gesellschaft zur Unterstützung der Krebsforschung und -therapie (GUK e.V.), the Research Program of the Comprehensive Cancer Center Göttingen (project 7-67-5343), Georg-August-University Göttingen, the German Ministry of Education and Research (BMBF) e:BIO project MetastaSys (0316173A) and e:Med project MyPathSem (031L0024) and the Excellence Cluster “Cells in Motion”. We thank L. Ries, M. Schaffrinski, M. Schulz, B. Celik and A. Westermann for their excellent technical assistance, and Gabriela Salinas (TAL: Microarray and Deep-Sequencing Core Facility Göttingen) for generation of sequencing data.

The authors declare no conflict of interest.

## AUTHOR CONTRIBUTIONS

K.M. and S.H. performed the in vitro experiments and analyzed the data. D.W. and M.S. compiled and analyzed high-throughput and patient data. H.T. and T.R. performed the electron microscopy. S.W. supervised and conceived the RPPA measurements. H.N. and A.J. designed, performed and analyzed the AFM and ECIS measurements. B.S., C.v.d.B. and T.P. provided patient samples. K.M., T.B., C.B. and A.B. conceived and designed the study and interpreted the data. K.M. wrote the manuscript.

## DATA AVAILABILITY STATEMENT

The RNA-Seq data of breast cancer brain metastasis and of MCF-7 pcDNA/pROR2 cells with/without siRNA treatment have been uploaded to the GEO repository under the identifiers GSE74383, GSE161864 and GSE161865. Uncropped scans of the western blots are provided in the source data. All other data supporting the findings of this study are available within the article and the supplemental information, or from the corresponding author upon reasonable request.

