## Supplementary for "WNT11 is a novel ligand for ROR2 in human breast cancer"

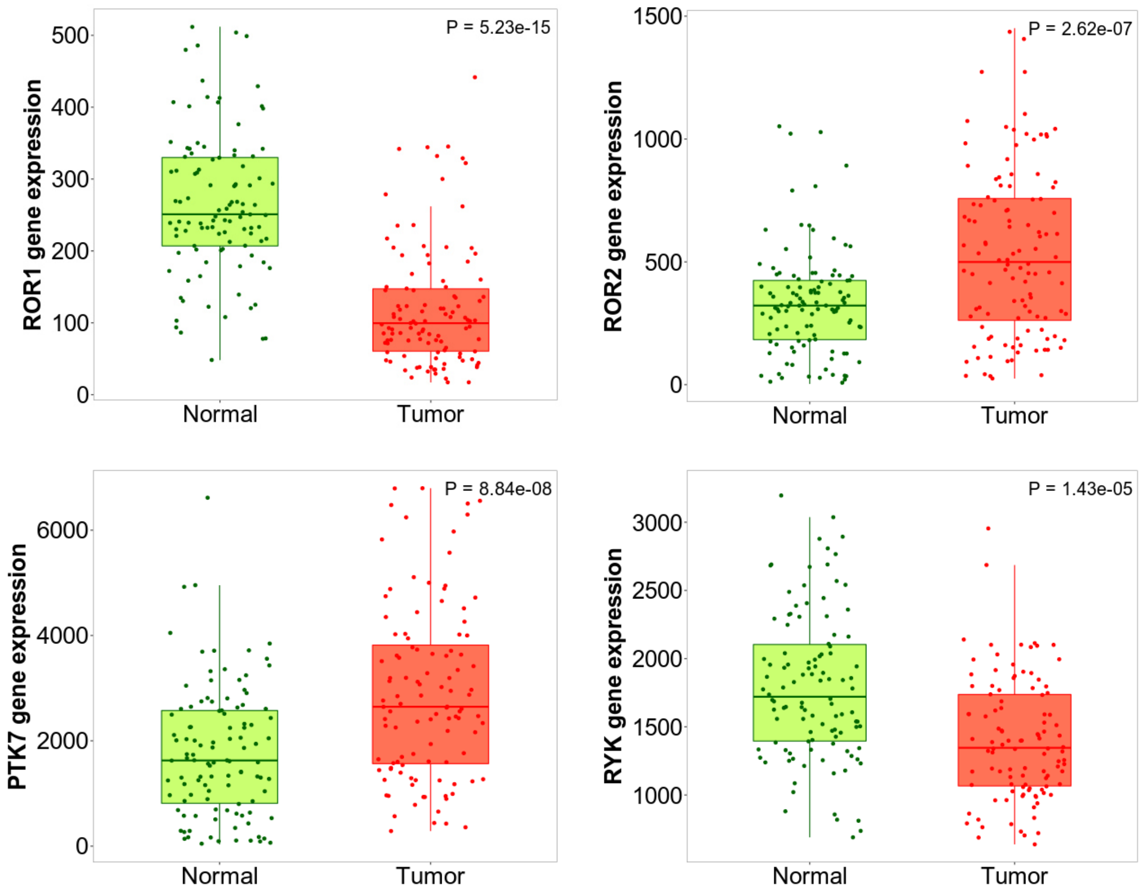


**Supplementary Fig. 1: ROR2 and PTK7 are overexpressed in human breast cancer.** Gene expression of the four non-canonical WNT co-receptors *ROR1, ROR2, PTK7* and *RYK* in normal breast (green) and matched breast cancer tissue (red) from the TNMplot database. Significance was calculated with a paired Wilcoxon statistical test.

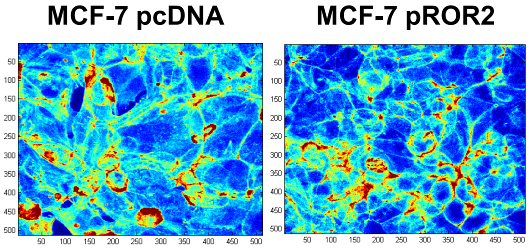


**Supplementary Fig. 2: The actin cytoskeleton in MCF-7 pROR2 cells.** Phalloidin staining of MCF-7 pcDNA and pROR2 cells confirming that the actin cytoskeleton is unchanged in pROR2 cells.

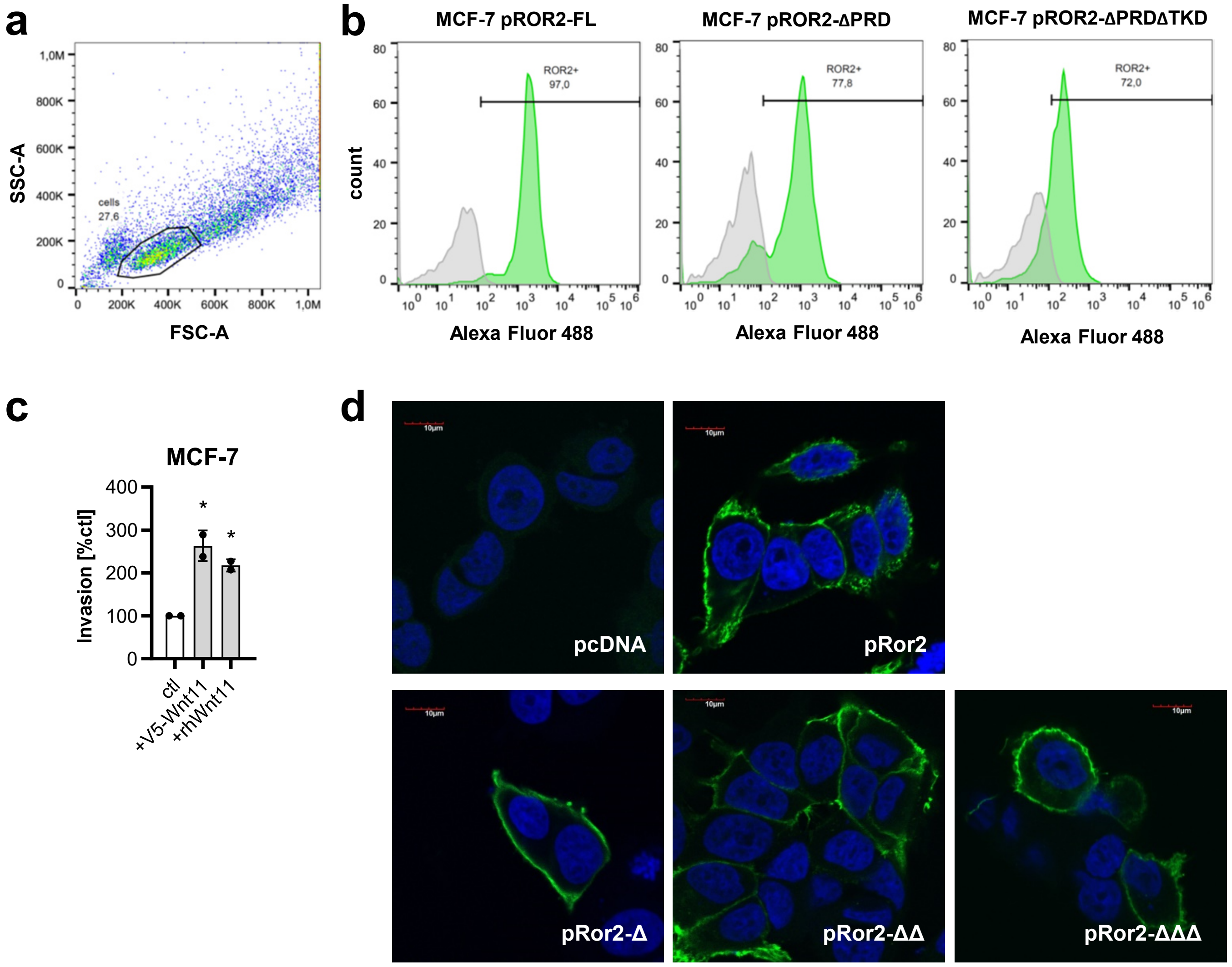


**Supplementary Fig. 3: Modulation of the WNT11 and ROR2 expression in MCF-7 cells.** ***a+b,*** MCF-7 cells stably transfected with either ROR2 full-length (pROR2-FL) or C-terminal deletion constructs were analyzed by flow cytometry for their ROR2 expression. Cell populations were gated based on FSC vs SSC plots (a) and representative histograms for the expression of ROR2 (green) in comparison to the isotype control (grey) in the gated cell population are shown (b). ***c,*** Invasion assay of MCF-7 cells transiently transfected with V5-tagged WNT11 (mean±SD, n=2, *p<0,001). ***d,*** Confocal microscopy: Immunofluorescence staining of MCF-7 cells transfected with the serial N-terminal ROR2 deletion constructs (blue: DAPI, green: ROR2). Scale bar: 10 µm

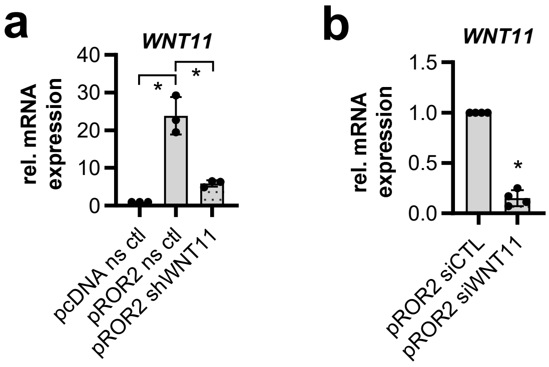


**Supplementary Fig. 4: Knockdown of WNT11 in MCF-7 pROR2 cells.** ***a***, qRT-PCR: *WNT11* knockdown was confirmed in MCF-7 cells stably overexpressing either a non-sense control (ns ctl) or a shRNA directed against *WNT11* (mean±SD, n=3, *p<0.01). ***b,*** *WNT11* expression was measured by qRT-PCR in MCF-7 pROR2 cells transiently transfected with a control siRNA (siCTL) or siRNA agsinst WNT11 (siWNT11) (mean±SD, n=4, *p<0.001).

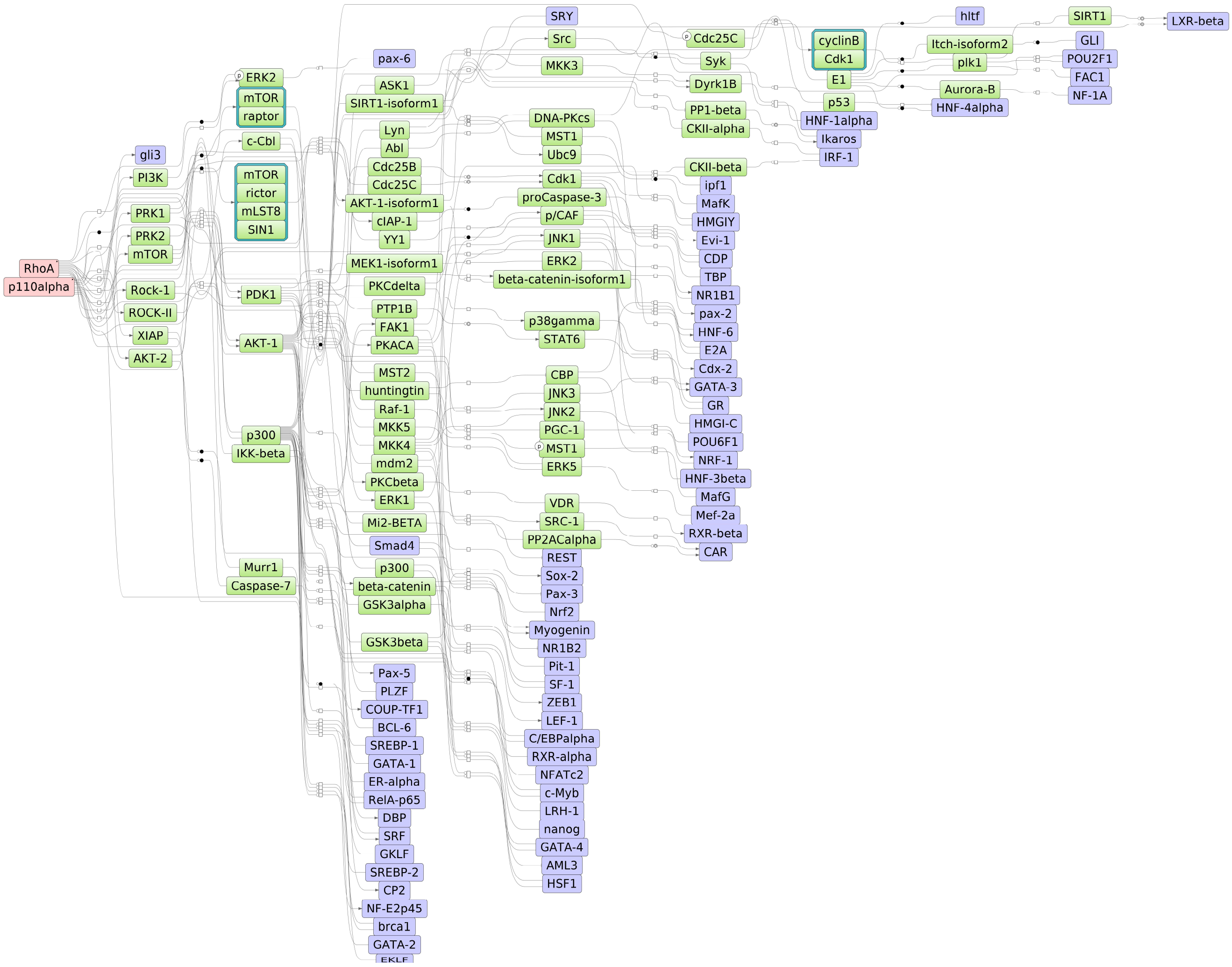


**Supplementary Fig. 5: PIK3CA and RHOA are master regulators of ROR2 signaling.** Master regulatory network based on RNA-Seq data of MCF-7 pcDNA vs pROR2 cells. Red: master regulators, purple: regulated transcription factors, green; connecting molecules.

**Supplementary Table 1: Antibodies used for RPPA**.

| Gene.Symbol | Target.Protein | UniProt.ID | Supplier | Antibody.ID | Antibody.Name | Phospho.Specificity | Host |
|---|---|---|---|---|---|---|---|
| *AKT1* | AKT1 | P31749 | CST | 4056 | Phospho-Akt (Thr308) (244F9) | T308 | rabbit |
| *AKT2* | AKT2 | P31751 | CST | 4056 | Phospho-Akt (Thr308) (244F9) | T309 | rabbit |
| *AKT3* | AKT3 | Q9Y243 | CST | 4056 | Phospho-Akt (Thr308) (244F9) | T305 | rabbit |
| *AKT1* | AKT1 | P31749 | CST | 4058 | Phospho-Akt (Ser473) (193H12) | S473 | rabbit |
| *AKT2* | AKT2 | P31751 | CST | 4058 | Phospho-Akt (Ser473) (193H12) | S474 | rabbit |
| *AKT3* | AKT3 | Q9Y243 | CST | 4058 | Phospho-Akt (Ser473) (193H12) | S472 | rabbit |
| *BRAF* | BRAF | P15056 | CST | 2696 | Phospho-B-Raf (Ser445) | S445 | rabbit |
| *CTNNB1* | CTNB1 | P35222 | CST | 9561 | Phospho-β-Catenin (Ser33/37/Thr41) | S33/S34/T41 | rabbit |
| *CTNNB1* | CTNB1 | P35222 | CST | 9567 | Phospho-β-Catenin (Ser675) | S675 | rabbit |
| *EGFR* | EGFR | P00533 | Millipore | 05-483 | Anti-phospho-EGFR (Tyr1173) Antibody, clone 9H2 | Y1173 | mouse |
| *EIF4EBP1* | 4EBP1 | Q13541 | CST | 2855 | Phospho-4E-BP1 (Thr37/46) (236B4) | T37/T46 | rabbit |
| *EIF4EBP2* | 4EBP2 | Q13542 | CST | 2855 | Phospho-4E-BP1 (Thr37/46) (236B4) | T37/T46 | rabbit |
| *EIF4EBP3* | 4EBP3 | O60516 | CST | 2855 | Phospho-4E-BP1 (Thr37/46) (236B4) | T23/T32 | rabbit |
| *ERBB2* | ERBB2 | P04626 | CST | 2243 | Phospho-HER2/ErbB2 (Tyr1221/1222)(6B12) | Y1221/Y1222 | rabbit |
| *ERBB3* | ERBB3 | P21860 | CST | 4791 | Phospho-HER3/ErbB3 (Tyr1289) (21D3) | Y1289 | rabbit |
| *GSK3A* | GSK3A | P49840 | Santa Cruz | 7291 | GSK-3α/β (0011-A) | n/a | mouse |
| *GSK3B* | GSK3B | P49841 | Santa Cruz | 7291 | GSK-3α/β (0011-A) | n/a | mouse |
| *GSK3A* | GSK3A | P49840 | CST | 9323 | Phospho-GSK-3β (Ser9) (5B3) | S21 | rabbit |
| *GSK3B* | GSK3B | P49841 | CST | 9323 | Phospho-GSK-3β (Ser9) (5B3) | S9 | rabbit |
| *GSK3A* | GSK3A | P49840 | CST | 9331 | Phospho-GSK-3α/β (Ser21/9) | S21 | rabbit |
| *GSK3B* | GSK3B | P49841 | CST | 9331 | Phospho-GSK-3α/β (Ser21/9) | S9 | rabbit |
| *GSK3A* | GSK3A | P49840 | CST | 9316 | Phospho-GSK-3α (Ser21) (36E9) | S21 | rabbit |
| *JUN* | c-Jun | P05412 | CST | 9164 | Phospho-c-Jun (Ser73) | S73 | rabbit |
| *JUND* | JunD | P17535 | CST | 9164 | Phospho-c-Jun (Ser73) | S100 | rabbit |
| *MAPK1* | ERK2 | P28482 | CST | 4370 | Phospho-p44/42 MAPK (Erk1/2) (Thr202/Tyr204) (D13.14.4E) | T185/Y187 | rabbit |
| *MAPK3* | ERK1 | P27361 | CST | 4370 | Phospho-p44/42 MAPK (Erk1/2) (Thr202/Tyr204) (D13.14.4E) | T202/Y204 | rabbit |
| *MAPK14* | p38MAPK | Q16539 | CST | 9211 | Phospho-p38 MAP Kinase (Thr180/Tyr182) | T180/Y182 | rabbit |
| *MEK1* | MEK1 | Q02750 | CST | 9154 | Phospho-MEK1/2 (Ser217/221) (41G9) | S218/S222 | rabbit |
| *MEK2* | MEK2 | P36507 | CST | 9154 | Phospho-MEK1/2 (Ser217/221) (41G9) | S222/S226 | rabbit |
| *MTOR* | mTOR | P42345 | CST | 2971 | Phospho-mTOR (Ser2448) | S2448 | rabbit |
| *PDPK1* | PDPK1 | O15530 | CST | 3061 | Phospho-PDK1 (Ser241) | S241 | rabbit |
| *PRKCA* | PKCα | P17252 | Abcam | 23513 | Anti-PKC alpha (phospho S657 + Y658) | S657/Y658 | rabbit |
| *PRKCA* | PKCα | P17252 | Millipore | 06-822 | Anti-phospho-PKCα (Ser657) | S657 | rabbit |
| *PRKCA* | PRKCA | P17252 | CST | 9375 | Phospho-PKCa/b II (Thr638/641) | T638 | rabbit |
| *PRKCB* | PRKCB | P05771 | CST | 9375 | Phospho-PKCa/b II (Thr638/641) | T641 | rabbit |
| *PRKCA* | PKCα | P17252 | CST | 9379 | Phospho-PKC (pan) (γ Thr514) | T497 | rabbit |
| *PRKCB* | PKCβI | P05771 | CST | 9379 | Phospho-PKC (pan) (γ Thr514) | T500 | rabbit |
| *PRKCG* | PKCγ | P05129 | CST | 9379 | Phospho-PKC (pan) (γ Thr514) | T514 | rabbit |
| *PRKCD* | PKCδ | Q05655 | CST | 9379 | Phospho-PKC (pan) (γ Thr514) | T507 | rabbit |
| *PRKCE* | PKCε | Q02156 | CST | 9379 | Phospho-PKC (pan) (γ Thr514) | T566 | rabbit |
| *PRKCH* | PKCη | P24723 | CST | 9379 | Phospho-PKC (pan) (γ Thr514) | T513 | rabbit |
| *PRKCQ* | PKCθ | Q04759 | CST | 9379 | Phospho-PKC (pan) (γ Thr514) | T538 | rabbit |
| *PTEN* | PTEN | P60484 | CST | 9549 | Phospho-PTEN (Ser380/Thr382/383) (44A7) | S380/T382/T383 | rabbit |
| *PTEN* | PTEN | P60484 | CST | 9551 | Phospho-PTEN (Ser380) | S380 | rabbit |
| *PTK2* | FAK | Q05397 | CST | 3283 | Phospho-FAK (Tyr397) | Y397 | rabbit |
| *RAF1* | cRAF | P04049 | CST | 9421 | Phospho-c-Raf (Ser259) | S259 | rabbit |
| *RAF1* | cRAF | P04049 | CST | 9431 | Phospho-c-Raf (Ser289/296/301) | S289/S296/S301 | rabbit |
| *RAF1* | cRAF | P04049 | CST | 9427 | Phospho-c-Raf (Ser338) (56A6) | S338 | rabbit |
| *RPS6* | RPS6 | P62753 | CST | 4858 | Phospho-S6 Ribosomal Protein (Ser235/236) (D57.2.2E) | S235/S236 | rabbit |
| *SRC* | Src | P12931 | CST | 2101 | Phospho-Src Family (Tyr416) | Y416 | rabbit |
| *LYN* | Lyn | P07948 | CST | 2101 | Phospho-Src Family (Tyr416) | Y379 | rabbit |
| *FYN* | Fyn | P06241 | CST | 2101 | Phospho-Src Family (Tyr416) | Y420 | rabbit |
| *LCK* | Lck | P06239 | CST | 2101 | Phospho-Src Family (Tyr416) | Y394 | rabbit |
| *YES1* | Yes | P07947 | CST | 2101 | Phospho-Src Family (Tyr416) | Y426 | rabbit |
| *HCK* | Hck | P08631 | CST | 2101 | Phospho-Src Family (Tyr416) | Y411 | rabbit |

**Supplementary Table 2: Primers used for quantitative real-time PCR**.

| primer | sequence forward primer (5'-3') | sequence reverse primer (5'-3') |
|---|---|---|
| hsWNT4_fw/rv150 | CTGAAGGAGAAGTTTGATGGTG | TGTCCTGCTCACAGAAGTC |
| hsWNT5A_fw/rv109 | AGGGCTCCTACGAGAGTGCT | GACACCCCATGGCACTTG |
| hsWNT6_fw/rv150 | GGAGCGTTTAAAGGACACTG | GATACTAACCTCACCCACCA |
| hsWNT11_fw/rv102 | CTCGGAACTCGTCTATCTG | GTTGGATGTCTTGTTGCAC |
| hsROR2_fw/rv143 | TTCTTCTTGGTTTGCATGTG | CTGATCTCTTTGAGTTTGGC |
| hsHPRT1_fw/rv89 | TAT GCT GAG GAT TTG GAA AGG | CAT CTC CTT CAT CAC ATC TCG |
| hsGNB2L1_fw/rv84 | AAC CCT ATC ATC GTC TCC T | CAA TGT GGT TGG TCT TCA G |

### Supplementary Table 3: Master regulator analysis of MCF-7 pcDNA siWNT11 versus pROR2 siCTL cells.

| Gene Symbol | Master molecule name | FDR |
|---|---|---|
| RHOA | RhoA(h) | 0,002 |
| RAD23A | Rad23A(h) | 0,002 |
| CUX1 | CDP(h) | 0,002 |
| UBE2K | HIP2(h) | 0,002 |
| AMER1 | FAM123B(h) | 0,003 |
| CREBBP | CBP(h) | 0,004 |
| NR3C1 | GR(h) | 0,005 |
| CEBPA | C/EBPalpha(h) | 0,005 |
| CHUK | IKK-alpha:IKK-beta{p}:(IKK-gamma)2 | 0,006 |
| IKBKB | IKK-alpha:IKK-beta{p}:(IKK-gamma)2 | 0,006 |
| IKBKG | IKK-alpha:IKK-beta{p}:(IKK-gamma)2 | 0,006 |
| RARA | NR1B1(h) | 0,006 |
| PIK3CA | p110alpha(h) | 0,007 |
| PKN2 | PRK2(h) | 0,009 |
| CASP6 | proCaspase-6(h) | 0,01 |
| PRSS1 | trypsin-1(h) | 0,01 |
| EP300 | p300(h) | 0,011 |
| NLRC4 | CLAN(h) | 0,011 |
| INS | insulin(h) | 0,011 |
| PHLPP1 | PHLPP(h) | 0,012 |
| PHLPP2 | phlpp2(h) | 0,012 |
| PIK3CD | p110delta(h) | 0,013 |
| MECOM | Evi-1(h) | 0,013 |
| RARB | NR1B2(h) | 0,013 |
| CASP1 | Caspase-1(h) | 0,014 |
| CAMK2A | CamKII(h) | 0,015 |
| CAMK2B | CamKII(h) | 0,015 |
| CAMK2D | CamKII(h) | 0,015 |
| CAMK2G | CamKII(h) | 0,015 |
| RAC3 | Rac3(h) | 0,015 |
| RHOG | RhoG(h) | 0,015 |
| REST | REST(h) | 0,015 |
| HNF4A | HNF-4alpha(h) | 0,015 |
| RAC2 | Rac2(h) | 0,016 |
| NEK2 | Nek2A(h) | 0,017 |
| TRIM32 | HT2A(h) | 0,017 |
| TGFBR2 | TGFbetaR-II(h) | 0,018 |
| GATA3 | GATA-3(h) | 0,019 |
| HSF1 | HSF1(h) | 0,019 |
| NANOG | nanog(h) | 0,02 |
| HTT | huntingtin(h) | 0,021 |
| ROCK2 | ROCK-II(h) | 0,022 |
| FURIN | PACE(h) | 0,022 |
| ESR1 | ER-alpha(h) | 0,022 |
| PDPK1 | PDK1-isoform2(h) | 0,023 |
| EGF | EGF:ErbB1{pY}:ErbB2{pY}:Src | 0,023 |
| EGFR | EGF:ErbB1{pY}:ErbB2{pY}:Src | 0,023 |
| ERBB2 | EGF:ErbB1{pY}:ErbB2{pY}:Src | 0,023 |
| SRC | EGF:ErbB1{pY}:ErbB2{pY}:Src | 0,023 |
| NFATC2 | NFATc2(h) | 0,025 |
| MYB | c-Myb(h) | 0,025 |
| CDC42 | Cdc42-isoform2(h) | 0,026 |
| PPP1CB | PP1-beta(h) | 0,027 |
| CDK1 | Cdk1(h) | 0,027 |
| CASP7 | proCaspase-7(h) | 0,029 |
| RELA | RelA-p65(h) | 0,029 |
| PRKCD | PKCdelta(h){pT507}{pS645}{pS664} | 0,03 |
| IGF1R | IGF-1R(h) | 0,03 |
| CASP2 | Caspase-2(h) | 0,031 |
| HMGA1 | HMGIY(h) | 0,031 |
| PTK2B | Pyk2-isoform1(h) | 0,032 |
| FRAT1 | Frat1(h) | 0,032 |
| PRKCG | PKCgamma(h) | 0,033 |
| SUMO1 | sumo1(h) | 0,033 |
| AKT1 | AKT(h){ub}n | 0,034 |
| AKT2 | AKT(h){ub}n | 0,034 |
| PRKCI | PKCiota(h) | 0,036 |
| DRD1 | D1(h) | 0,036 |
| PDK1 | pyruvate dehydrogenase (lipoamide) kinase isozyme 1, mitochondrial(h) | 0,037 |
| BAX | Bax(h) | 0,038 |
| CASP10 | Caspase-10(h) | 0,04 |
| SENP5 | Senp5-isoform1(h) | 0,04 |
| DLG4 | PSD-95(h) | 0,04 |
| DUSP22 | JKAP(h) | 0,042 |
| MTOR | mTOR(h):rictor(h) | 0,042 |
| RICTOR | mTOR(h):rictor(h) | 0,042 |
| EPHB2 | EPHB2(h) | 0,042 |
| SGK1 | SGK-1(h){pT256} | 0,042 |
| FAS | Fas(h) | 0,044 |
| BRCA1 | brca1(h) | 0,045 |
| SAE1 | Aos1(h):SAE2-isoform1(h) | 0,045 |
| UBA2 | Aos1(h):SAE2-isoform1(h) | 0,045 |
| PPP1R9B | neurabin-II(h) | 0,047 |
| MUL1 | MAPL(h) | 0,048 |
| SQSTM1 | p62(h) | 0,049 |

**Supplementary Table 4: Master regulator analysis of MCF-7 pcDNA siWNT11 versus pROR2 siWNT11 cells.**

| Gene Symbol | Master molecule name | FDR |
|---|---|---|
| PRSS1 | trypsin-1(h) | 0,001 |
| RHOA | RhoA(h) | 0,002 |
| CUX1 | CDP(h) | 0,003 |
| IGF1R | IGF-1R(h) | 0,003 |
| EGF | EGF:ErbB1{pY}:ErbB2{pY}:Src | 0,008 |
| EGFR | EGF:ErbB1{pY}:ErbB2{pY}:Src | 0,008 |
| ERBB2 | EGF:ErbB1{pY}:ErbB2{pY}:Src | 0,008 |
| SRC | EGF:ErbB1{pY}:ErbB2{pY}:Src | 0,008 |
| YY1 | YY1(h) | 0,01 |
| ESR1 | ER-alpha(h) | 0,01 |
| DUSP22 | JKAP(h) | 0,011 |
| NR3C1 | GR(h) | 0,011 |
| CEBPA | C/EBPalpha(h) | 0,011 |
| PPP1CB | PP1-beta(h) | 0,012 |
| PPP1R9B | neurabin-II(h) | 0,012 |
| MYB | c-Myb(h) | 0,014 |
| MECOM | Evi-1(h) | 0,015 |
| PRKCD | PKCdelta(h),PKCdelta(h){pT507}{pS645}{pS664} | 0,016 |
| RARA | NR1B1(h) | 0,018 |
| GATA3 | GATA-3(h) | 0,02 |
| RARB | NR1B2(h) | 0,02 |
| CHUK | IKK-alpha:IKK-beta{p}:(IKK-gamma)2 | 0,021 |
| IKBKB | IKK-alpha:IKK-beta{p}:(IKK-gamma)2 | 0,021 |
| IKBKG | IKK-alpha:IKK-beta{p}:(IKK-gamma)2 | 0,021 |
| INS | insulin(h) | 0,021 |
| NANOG | nanog(h) | 0,021 |
| LMTK2 | KPI-2(h) | 0,022 |
| NFATC2 | NFATc2(h) | 0,022 |
| KRAS | K-Ras(h) | 0,023 |
| PTK2B | Pyk2-isoform1(h) | 0,023 |
| NEK2 | Nek2A(h),Nek2A(h){p} | 0,024 |
| CASP2 | proCaspase-2(h) | 0,024 |
| GABPA | GABP-alpha(h) | 0,024 |
| HSF1 | HSF1(h) | 0,024 |
| DRD1 | D1(h) | 0,026 |
| GABPB1 | GABP-beta(h) | 0,026 |
| RAD23A | Rad23A(h) | 0,027 |
| AMER1 | FAM123B(h) | 0,027 |
| HMGA1 | HMGIY(h) | 0,027 |
| PPP1CA | PP1-alpha(h),PP1-alpha1(h),PP1-alpha2(h) | 0,03 |
| RAC3 | Rac3(h) | 0,034 |
| RHOG | RhoG(h) | 0,034 |
| PPP1CC | PP1-gamma1(h) | 0,035 |
| ICMT | ICMT(h) | 0,035 |
| BAIAP2 | IRSp53(h) | 0,036 |
| IRS4 | IRS-4(h) | 0,037 |
| DUSP10 | MKP-5-isoform1(h) | 0,038 |
| ARHGDIA | RhoGDI-1(h) | 0,038 |
| HTT | huntingtin(h) | 0,039 |
| RAC2 | Rac2(h) | 0,039 |
| RELA | RelA-p65(h) | 0,042 |
| CDC42 | Cdc42-isoform1(h),Cdc42-isoform2(h) | 0,043 |
| FURIN | PACE(h) | 0,0435 |
| CAMK2A | CamKII(h) | 0,045 |
| CAMK2B | CamKII(h) | 0,045 |
| CAMK2D | CamKII(h) | 0,045 |
| CAMK2G | CamKII(h) | 0,045 |
| ROCK2 | ROCK-II(h) | 0,047 |
| FAS | Fas(h) | 0,047 |
